# KIF1A neurodegenerative disease mutations modulate motor motility and force generation

**DOI:** 10.1101/2023.06.09.544228

**Authors:** Dipeshwari J. Shewale, Pushpanjali Soppina, Virupakshi Soppina

**Affiliations:** Discipline of Biological Engineering, Indian Institute of Technology Gandhinagar, Gandhinagar, Gujarat 382355, India; Department of Biotechnology and Bioinformatics, Sambalpur University, Sambalpur, Orissa 768019, India

**Keywords:** KIF1A, Kinesin-3, Microtubule, Neurodegeneration, Superprocessive

## Abstract

KIF1A is involved in fast axonal transport of synaptic vesicle precursor, neurofilament, and dense-core vesicle, essential for neuronal development and maintenance. Several point mutations in the KIF1A motor domain have been identified in patients with various motor neuron diseases. Recent studies have shown that these mutations affected the motor and cargo localization in cultured hippocampal and *C.elegans* neurons. However, a detailed analysis of these mutations on KIF1A motility, force generation, and cargo transport is largely unexplored. Here, we have analyzed the effect of 16 point mutations and showed that these mutations significantly decreased the motor velocity and landing rates compared to wild-type motors. Except for A255V, mutations V144F, V220I, and E233D mildly affected motor mechanical outputs. S58L, A202P, R216P, R216H, L249Q, T312M, and R316W mutants exhibited drastic impairments in the motility properties, force generation, and cargo transport. Notably, T46M, T99M, G102D, S215R, and E253K mutants showed strong microtubule binding, resulting in complete disruption of cargo transport. Our study provides the first comprehensive demonstration of KIF1A disease mutations at the molecular level. The observed changes in motility properties and cargo transport align with the severity of disease phenotype observed in KIF1A-associated neurological disorders)

## Introduction

Neurons are highly polarized cell types with axons, long cable-like structures wiring the entire nervous system from the cell body. Most proteins required for neuronal growth, maintenance, and signaling are synthesized in the cell body and delivered to the synapse via a long-distance microtubule-based transport system. Axonal microtubules are arranged in parallel bundles, with their plus-ends extending away from the cell body and facing the synapse. Microtubule-based kinesin and dynein motor proteins walk along these microtubule bundles in opposite directions while transporting the cargo. Kinesins transport towards the plus-end while dynein to the minus-end of the microtubule. KIF1A, a novel neuron-specific motor, belongs to the kinesin-3 family involved in fast axonal transport of synaptic vesicle proteins (SVPs), neurofilament, and dense-core vesicles essential for the development, maintenance, and neuronal signaling (Hall and Hedgecock, 1991; Hirokawa et al., 2009; Lo et al., 2011; Okada et al., 1995; Vale, 2003; Verhey et al., 2011). KIF1A interacts with a scaffold protein liprin-α and transports vesicles containing post-synaptic proteins, such as GRIP, GIT1, and AMPA receptors, which play an important role in synaptic plasticity and transmission as well as learning and memory (Shin et al., 2003). Since then, numerous studies have been published on the strong association of KIF1A with several neuronal cargoes, such as TrkA neurotrophin receptors and β-secretase-1 (BACE-1), crucial for cell survival of dorsal root ganglia and superior cervical ganglia (Gabrych et al., 2019; Siddiqui and Straube, 2017).

Akin to other kinesins, KIF1A has a conserved motor domain (MD) with both catalytic activity and microtubule-binding ability. MD is followed by a flexible linker to a dimerization domain neck coil (NC), coiled-coil1(CC1), a family-specific fork-head associated (FHA), coiled-coil2 (CC2), and coiled-coil3 (CC3) domains (Scarabelli et al., 2015; Soppina et al., 2022d; Soppina et al., 2014; Soppina and Verhey, 2014). It also has scaffold binding and C-terminal lipid-binding pleckstrin homology (PH) region. Additionally, Kloop, a family-specific positively charged insert in the loop12 rich in lysine residues, forms electrostatic interactions with the negatively charged glutamine-rich C-terminal of tubulin, thereby influencing microtubule affinity of KIF1A (Nitta et al., 2004; Okada and Hirokawa, 1999; Okada and Hirokawa, 2000; Soppina and Verhey, 2014).

Unlike kinesin-1, KIF1A exhibits inherently fast and superprocessive motility with high affinity for the microtubule (Soppina et al., 2022a; Soppina et al., 2022c; Soppina et al., 2014; Soppina and Verhey, 2014). Mutations in the KIF1A motor are observed in multiple neurodegenerative diseases like Hereditary sensory autonomic neuropathy type-2 (HSAN II) and Hereditary spastic paraplegia (HSP), particularly SPG30. HSP is characterized by lower extremity spasticity and weakness in its pure form and can include symptoms like cognitive impairment and ataxia in its complicated forms (Klebe et al., 2012; Ohba et al., 2015; Okamoto et al., 2014). It is known to have both autosomal dominant and recessive inheritance patterns based on the mutation. Several de novo mutations have been found in the KIF1A motor domain leading to loss of motility or function of the motor. Mutations like T99M and E253K occur with higher frequency than others (Esmaeeli Nieh et al., 2015; Lee et al., 2015). Lee et al. reported 11 heterozygous de novo mutations in KIF1A, and transfection studies of 8 mutants (T99M, A202P, S215R, R216P, E253K, and R350G) in cultured hippocampal neurons showed drastically reduced tip accumulation as compared to A255V and wild-type (WT) motor (Lee et al., 2015). A similar study with T99M and T312M mutants displayed their accumulation in the cell body rather than the tip of an axon, as observed with WT KIF1A (Hamdan et al., 2011). The autosomal dominant mutation T258M was evaluated in complicated HSP, and it showed decreased peripheral accumulation of the mutant, which was not rescued by the WT motor. This suggested that it might be acting in a dominant negative manner (Cheon et al., 2017). Microtubule gliding analysis of five mutants depicted that V220I and A255V showed gliding velocity similar to WT motor, whereas T99M, R216C, and E253K mutants did not show any gliding events (Esmaeeli Nieh et al., 2015). The motility studies by Chiba et al. suggested that V8M and A255V mutations hyperactivated KIF1A motor due to their increased landing rates and resulted in higher axonal transport of SVPs and their abnormal accumulation (Chiba et al., 2019). In contrast, A255V mutation showed significantly lower run length and velocity than WT motor in the motility assay (Guedes-Dias et al., 2019). Although KIF1A disease mutations were investigated in cultured hippocampal neurons, there is a lack of knowledge about the effect of these mutations on motor motility and cargo transport. In the present study, we have performed a comprehensive analysis of KIF1A neurodegenerative disease mutations and demonstrated that these mutations cause a defect in motor motility properties, force generation, and cargo transport. This study will help us understand the functioning of KIF1A mutants at the molecular level and may help overcome the defective neuronal functions in KIF1A-related diseases in the near future. Our findings provide a fundamental understanding of the mechanisms underlying KIF1A-related diseases, thereby paving the way for potential therapeutic interventions targeting these mutations.

## Results

### Kinesin-3 motors accumulate at the cell periphery due to their long run length

Dimeric constitutively active kinesin-3 motors [KIF1A(1-393LZ), KIF13A(1-411ΔP), KIF13B(1-412ΔP), and KIF16B(1-400)] exhibit fast velocity and superprocessive motility (10-fold longer processive than kinesin-1) (Soppina et al., 2014) with strong affinity for microtubule than kinesin-1 (Soppina and Verhey, 2014). Interestingly, the expression of constitutively active kinesin-3 motors in COS-7 cells remarkably localized to the cell periphery (**Fig. 1A**). For clarity and relevance, we have shown KIF1A(1-393LZ) results alone. On the contrary, expression of constitutively active kinesin-1 motor, KHC(1-560), showed uniform cytoplasmic distribution with no significant peripheral localization (**Fig. 1A**).

**Figure 1:**
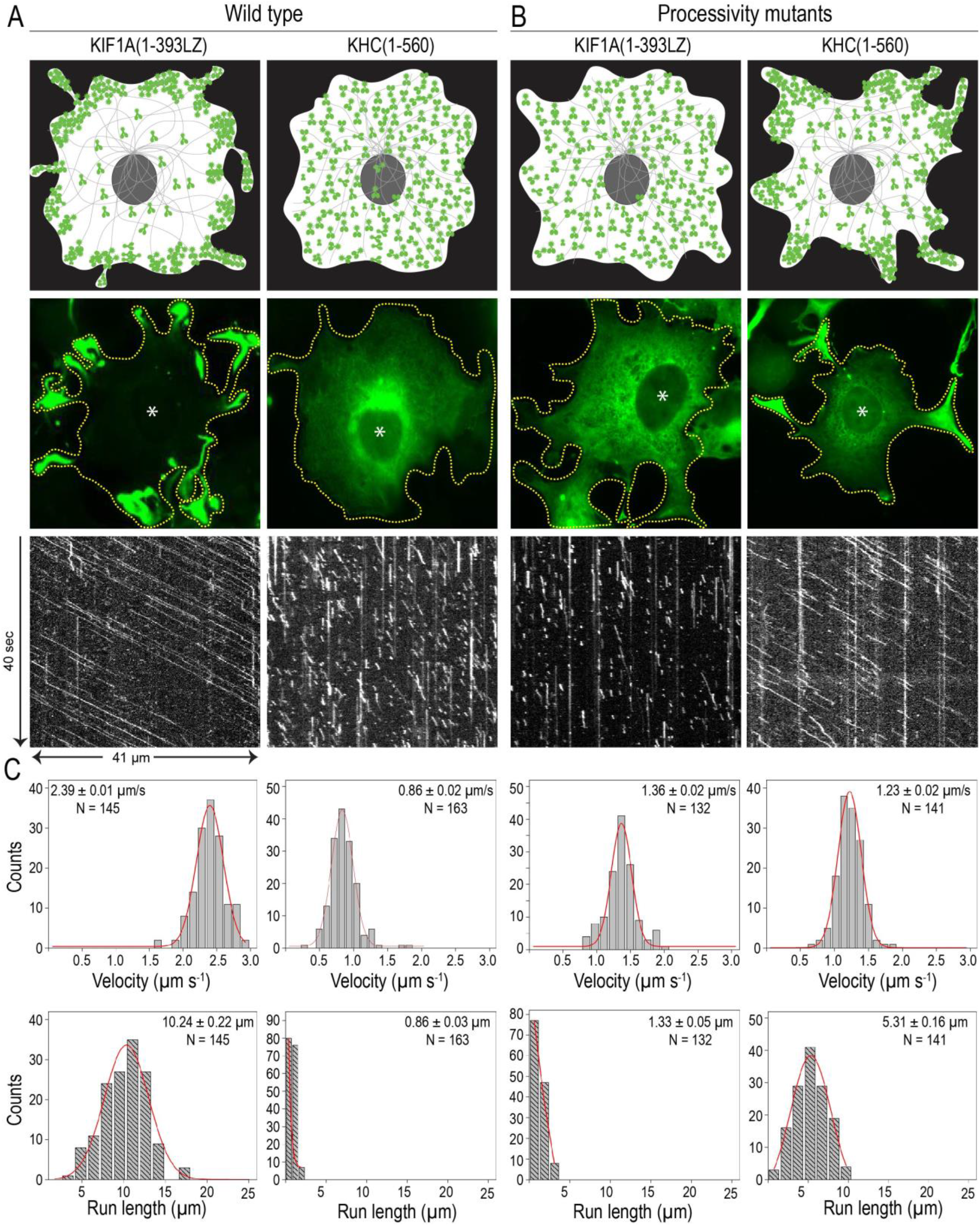
KIF1A processivity leads to its localization at the cell periphery. Cartoon representation of cellular localization of motors, expression of motors in COS-7 cells, and kymograph from single molecule motility assay of **A.** constitutively active, wild-type KIF1A(1-393LZ) and KHC(1-560) motor with peripheral and cytoplasmic distribution respectively and **B.** processivity mutants KIF1A(1-393LZ-R346M) and KHC(1-560-A155R/H157R/M346R) with localization in contrast with their wild-type motors. **C.** Histogram fitted to Gaussian distribution for velocity and run-length after tracking multiple events (displayed in the right corner of the plots by N) along with velocity ± SEM.

One possibility would be that kinesin-3 motors preferentially bind to GTP tubulin cap located at the growing plus-ends of the microtubules located at the cell periphery; consequently, motors localized to the cell periphery. Indeed, KIF7, a kinesin-4 family member, has been shown to preferentially bind to GTP-tubulin at microtubule ends and localize to the ciliary tip (Jiang et al., 2019). Microtubules exhibit dynamic instability with an average microtubule growth rate of 0.2–0.4 µms^-1^, significantly slower than the measured motor velocity and shrinks rapidly (Zwetsloot et al., 2018). Therefore, this mechanism cannot explain the unidirectional inherently fast plus-end superprocessive motility of the kinesin-3 motors. An alternative possibility is that most kinesin-3 motors contain a lipid-binding domain in their C-terminal tail region that could interact with the plasma membrane at the cell periphery (Klopfenstein et al., 2002; Klopfenstein and Vale, 2004). This mechanism also seems unlikely to explain the peripheral accumulation of kinesin-3 motors because the motors used in the assay are truncated and contain a catalytic motor domain, a short neck linker, and a neck coil domain for motor dimerization.

Next, we wondered whether the prominent accumulation of truncated dimeric kinesin-3 motors to the cell periphery is likely due to their inherent superprocessive motility along the microtubules. As microtubules in COS-7 cells are arranged radially with plus-ends extending to the cell periphery, kinesin-3 motors travel along these microtubules and reach the cell periphery before their detachment. If motors detach, their higher microtubule affinity enables them to rebind quickly to the microtubules and walk to the microtubule plus-ends, accumulating at the cell periphery. Conversely, kinesin-1 motors are short processive, traveling a short distance before detaching from the microtubule. Once detached, they need much longer to rebind to microtubules due to their low affinity for the microtubule, rendering the motors largely cytoplasmic.

To test this hypothesis, we introduced a point mutation, R346M, into KIF1A(1-393LZ) motor domain to make it less processive. R346M is a conserved residue located at the motor-microtubule interface and is critical for establishing strong electrostatic interactions with microtubules while propagating processive motility (Scarabelli et al., 2015; Uchimura et al., 2010). Similarly, we generated a kinesin-1 triple mutant by mutating residues A155R and H157R in Loop8 and M346R in KHC(1-560) motor, making the motor longer processive (Scarabelli et al., 2015). To examine the effect of these mutations on motor processivity and their cellular localization, we expressed these mutants in COS-7 cells and assessed their cellular distribution. Cells expressing KIF1A(1-393LZ-R346M) showed diffuse cytoplasmic distribution (**Fig. 1B**) as opposed to the peripheral accumulation of WT KIF1A(1-393LZ) motor. Interestingly, expression of kinesin-1 triple mutant, KHC(1-560-A155R/H157R/M346R), showed significant cell peripheral accumulation, which is comparable to WT KIF1A(1-393LZ) motor (**Fig. 1**, A and B).

Next, we assessed the motility properties of these mutants by TIRF (Total Internal Reflection Fluorescence) microscopy-based single-molecule motility assays. Cell lysates containing WT or mutant motors tagged with three tandem monomeric citrine fluorescent proteins (3xmCit) were introduced into flow chambers adsorbed with taxol-stabilized microtubules. Motility analysis of KIF1A(1-393LZ) exhibited long processive motility along the microtubule with an average velocity of 2.39 µms^-1^ and run-length of 10.24 µm (**Fig. 1A**), consistent with previous studies (Scarabelli et al., 2015; Soppina and Verhey, 2014). However, the mutant motor, KIF1A(1-393LZ-R346M), showed short processive runs with an average velocity of 1.36 µms^-1^ and run-length of 1.33 µm (**Fig. 1B**). On the other hand, motility analysis of WT KHC(1-560) showed typical short processive runs with an average velocity of 0.86 µms^-1^ and run-length 0.86 µm (**Fig. 1A**), as shown previously. Intriguingly, the motility assay with mutant kinesin-1 displayed longer processive motion along the microtubule with an average velocity of 1.23 µms^-1^ and run-length of 5.31 µm, which is 5-fold higher than WT kinesin-1 motor (**Fig. 1B**). Together, these results demonstrate that peripheral accumulation of kinesin-3 motors in COS-7 cells is indeed due to their long processive motility and is independent of motor velocity.

### KIF1A disease mutants exhibit distinct cellular localization

KIF1A, a founding member of the kinesin-3 family, is responsible for anterograde transport of diverse neuronal cargoes, critical for pre-and post-synaptic assembly, neuronal function and survival (Gabrych et al., 2019; Niwa et al., 2008; Okada et al., 1995; Otsuka et al., 1991). A large number of autosomal dominant or recessive mutations in the motor domain of KIF1A have been correlated with a wide spectrum of neurodegenerative disorders like Hereditary sensory autonomic neuropathy type-2 (HSAN II) and Hereditary spastic paraplegia (HSP), mainly SPG30 (Cheon et al., 2017; Citterio et al., 2015; Erlich et al., 2011; Esmaeeli Nieh et al., 2015; Nicita et al., 2021; Okamoto et al., 2014; Riviere et al., 2011). Kinesin motors utilize chemomechanical coupling of the ATP hydrolysis to step processively along the microtubule. Any alternations in the ATPase cycle or microtubule-binding due to mutations in the motor domain disrupt motor function, consequently, intracellular cargo transport. However, recent work from Chiba et al. demonstrated that some disease mutations hyperactivate KIF1A and increase the anterograde transport of synaptic vesicles (Chiba et al., 2019), which can also lead to improper cargo delivery.

To investigate the effect of neurodegenerative disease mutations in the KIF1A motor domain on motor’s motility properties and its intracellular localization, we incorporated each of these mutations into the constitutively active, dimeric motor, KIF1A (1-393LZ) (**Table 1**). To rapidly assess their cellular distribution, we employed the above COS-7 cell-based assay as described in **Fig. 1A**. Expression of individual mutants in COS-7 cells exhibited distinct cellular distribution (**Fig. 2**). WT KIF1A(1-393LZ) displayed a strong peripheral accumulation of motors due to their fast and superprocessive motility (**Fig. 2**). Based on the severity of the mutation, the assay clearly showed three different phenotypes on expression of KIF1A mutants in COS-7 cells 1) Peripheral: Mutations that showed a little-to-no effect on the motility properties of KIF1A showed significant peripheral accumulation of motors, 2) Cytoplasmic: Mutations that showed a drastic effect on KIF1A motility properties revealed uniform cytoplasmic distribution and 3) Microtubule bound: Mutations that caused KIF1A motor to bind microtubule, exhibiting microtubule decoration by the motors. Expression of V144F, V220I, E233D, and A255V mutants showed peripheral accumulation (hereinafter referred to as peripheral mutants) comparable to that of WT KIF1A(1-393LZ) (**Fig. 2; Fig. S1**), suggesting that these mutations likely have a mild effect on motor’s mechanochemical output.

**Figure 2:**
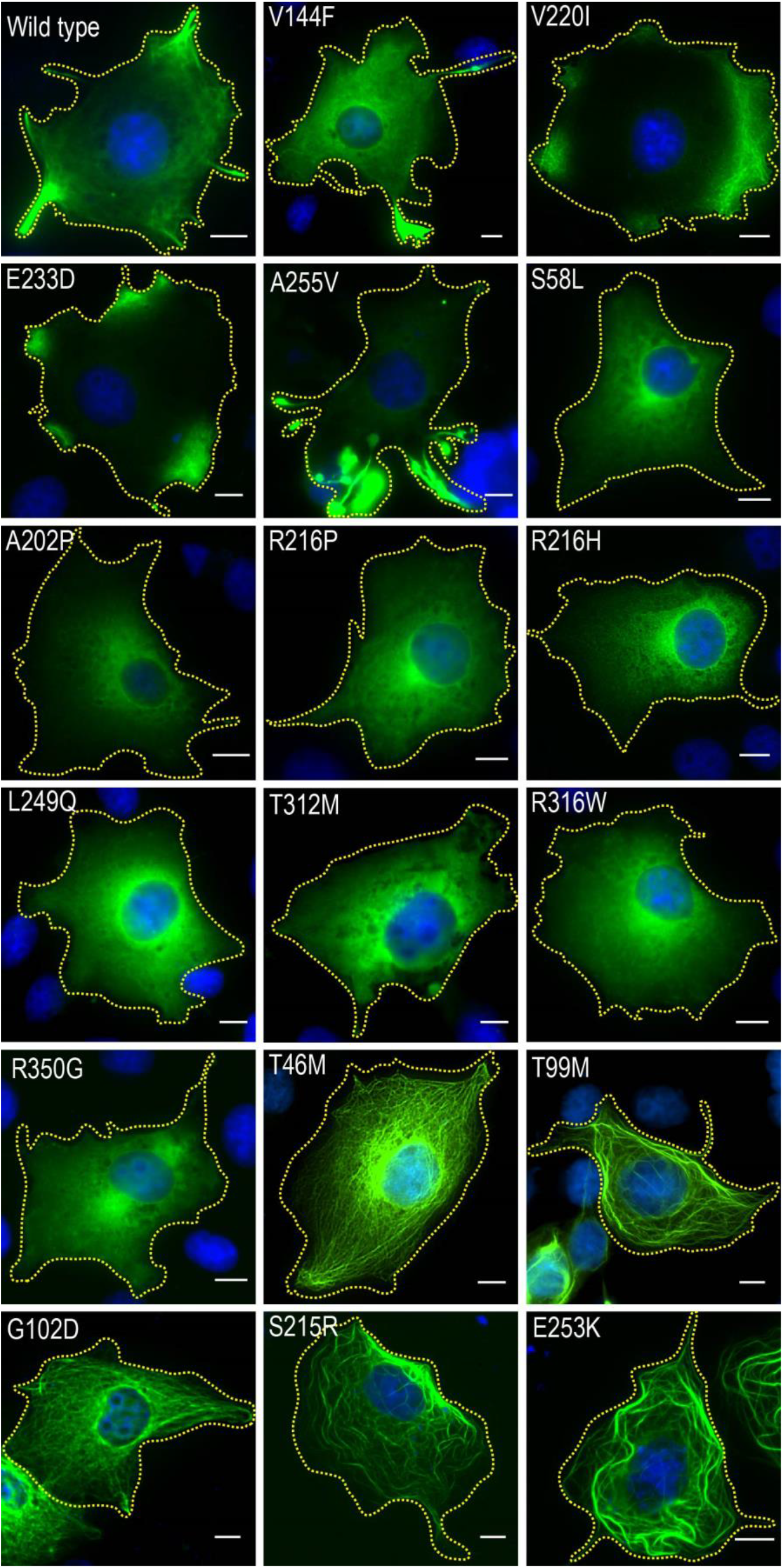
Distinct localization of KIF1A disease mutants in COS-7 cells. KIF1A(1-393LZ)-3XmCitrine motors with 17 point mutations were individually expressed in COS-7 cells and their localization was observed to be peripheral (V144F, V220I, E233D and A255V), diffused cytoplasmic (S58L, A202P, R216P, R216H, L249Q, R316W, and R350G) or microtubule bound (T46M, T99M, G102D, S215R and E253K). A representative merged image for each mutant is displayed (The green channel represents the motor; the blue channel shows DAPI stained nucleus). Scale Bar: 10µm.

**Table 1:** Consolidated phenotype of KIF1A WT and mutant motors in cell-based assays

| **Sr. No** | **KIF1A (1-393LZ)** | **COS-7 assay** | **CAD assay** | **Peroxisome assay** | **GTS assay** | **SMMA** |
| --- | --- | --- | --- | --- | --- | --- |
| 1 | WT | Periphery | Tip accumulation | Peripheral | Peripheral | Motile |
| 2 | V144F | Periphery | Tip accumulation | Partial | Partial | Motile |
| 3 | V220I | Periphery | Tip accumulation | Peripheral | Partial | Motile |
| 4 | E233D | Periphery | Tip accumulation | Peripheral & Partial | Partial | Motile |
| 5 | A255V | Periphery | Tip accumulation | Peripheral & Partial | Partial | Binds and Jiggles |
| 6 | S58L | Cytoplasmic | Cytoplasmic & Tip | Partial | Perinuclear | Moderately processive |
| 7 | A202P | Cytoplasmic | Cytoplasmic & Tip | Partial | Partial | Low processive |
| 8 | R216P | Cytoplasmic | Cytoplasmic & Tip | Tight | Tight | Low processive |
| 9 | R216H | Cytoplasmic | Cytoplasmic | Tight | Tight | Binds but not motile |
| 10 | L249Q | Cytoplasmic | Cytoplasmic | Perinuclear & Tight | Tight | Low processive |
| 11 | T312M | Cytoplasmic | Cytoplasmic | Perinuclear | Tight | Low processive |
| 12 | R316W | Cytoplasmic | Cytoplasmic & Tip | Partial | Partial | Binds and Jiggles |
| 13 | T46M | MT bound | MT bound | Perinuclear & MT bound | Perinuclear & MT bound | Non-motile |
| 14 | T99M | MT bound | MT bound | Perinuclear & MT bound | Perinuclear & MT bound | Non-motile |
| 15 | G102D | MT bound | MT bound | Perinuclear & MT bound | Perinuclear & MT bound | Non-motile |
| 16 | S215R | MT bound | MT bound | Perinuclear & MT bound | Perinuclear & MT bound | Non-motile |
| 17 | E253K | MT bound | MT bound | Perinuclear & MT bound | Perinuclear & MT bound | Non-motile |
| 18 | R350G | Cytoplasmic (less expression) | NA | NA |  | NA |

Similarly, S58L, A202P, R216P, R216H, L249Q, T312M, R316W, and R350G mutants showed diffused cytoplasmic localization (from now on referred to as cytoplasmic mutants) (**Fig. 2; Fig. S2**), suggesting that these mutations have strong and direct implications on motor’s processive motility. Therefore, these mutants presumably generate short processive motility similar to kinesin-1 or lose their ability to interact with microtubules, thus showing diffuse cytoplasmic distribution. Unlike other mutations, T46M, T99M, G102D, S215R, and E253K mutants did not show peripheral or cytoplasmic phenotype; instead, they showed uniform microtubule decoration (hereinafter referred to as microtubule bound) (**Fig. 2; Fig. S3**), rendering the motors immobile on the microtubule. Since previous *in vivo* expression studies were carried out in cultured hippocampal neurons and imaged at a lower resolution, the microtubule-binding phenotype of these mutants was not observed (Cheon et al., 2017; Lee et al., 2015).

### KIF1A disease mutants cause defects in neuronal growth and differentiation

KIF1A plays an indispensable role in the axonal transport of many presynaptic components to synapses and mutations that disrupt motor function cause defects in neuronal growth, differentiation, and function. As these mutations are implicated in neurodegenerative diseases, we wanted to examine their effect on neuronal structure and differentiation. Additionally, the COS-7 assay does not reveal whether mutant motors displayed diffused cytoplasmic localization due to low processivity or inability to generate microtubule-based motility. To probe this, we employed CAD cell assay, in which neuronal differentiation can be induced and maintained by serum starvation (Qi et al., 1997). Upon serum deprivation, CAD cells differentiate to form neuronal-like processes, where microtubules are arranged in an isopolar orientation with their minus ends facing towards the cell body and plus ends toward neurite tips. This enables the characteristic accumulation of constitutively active kinesin-3 motors to the neurite tips, whereas inactive motors remain in the cell body (Jacobson et al., 2006; Lee et al., 2004; Nakata and Hirokawa, 2003; Patel et al., 2021; Soppina et al., 2014).

As expected, CAD cell expression of peripheral mutants, V144F, V220I E233D and A255V, displayed strong neurite tip accumulation comparable to WT motors (**Fig. 3, A and B**). Next, we expressed cytoplasmic mutants in CAD cells. Although these mutants showed cytoplasmic distribution in COS-7 cells, only S58L, A202P, T312M and R316W mutants accumulated at the neurite tips, albeit with significantly less efficiency than WT motors (**Fig. 3, A and B**), hinting that these mutants bind to microtubules and exhibit processive motility. The remaining cytoplasmic mutants R216P, R216H and L249Q failed to show accumulation at the neurite tip and remain localized in the cell body (**Fig. 3, A and B**), indicating that these mutants lost their ability to interact with microtubules and consequently failed to generate processive motility.

**Figure 3:**
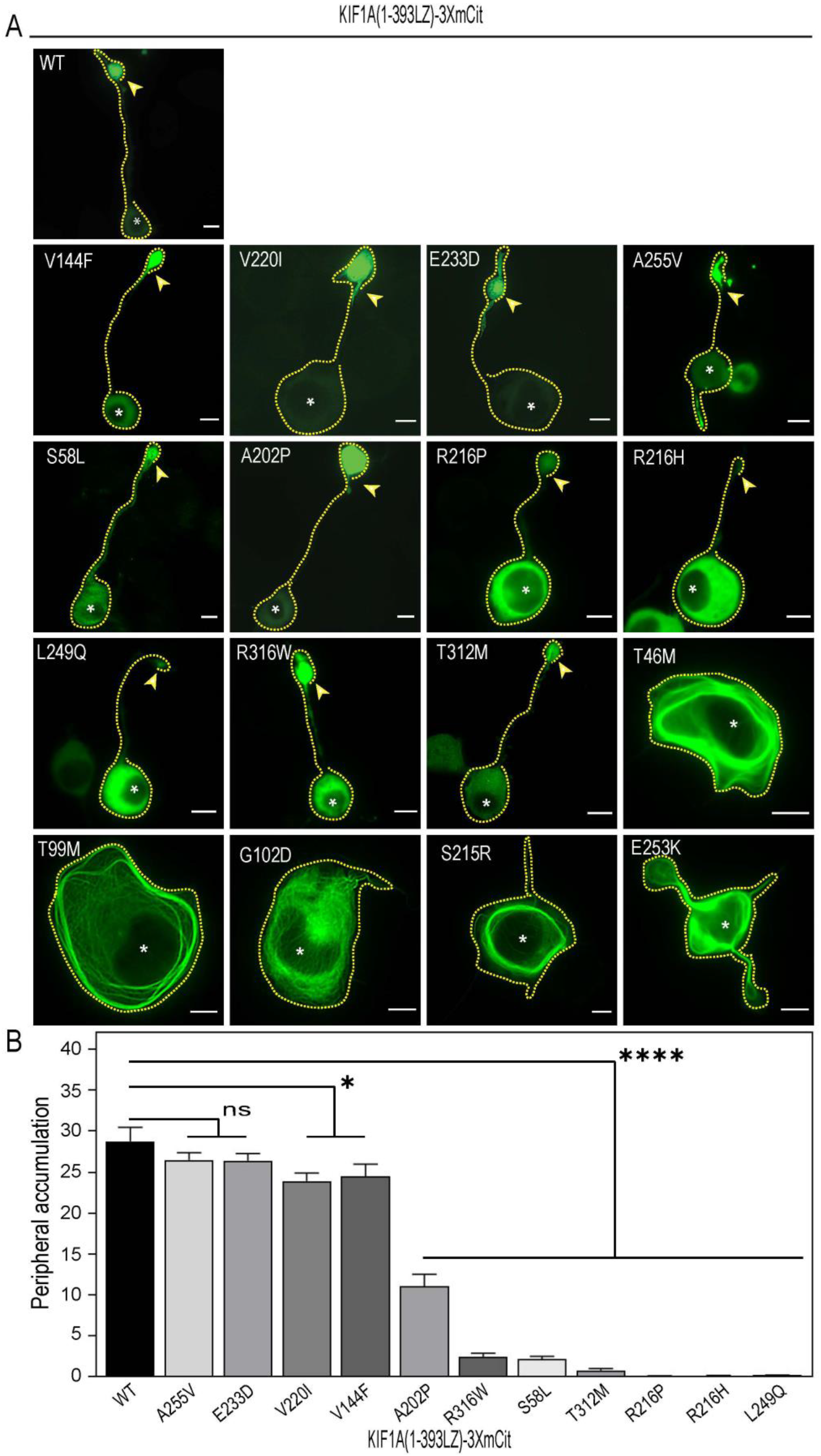
Effect of KIF1A disease mutants on tip accumulation and differentiation in CAD cells. Disease mutants with 16 individual mutations were transfected in CAD cells, and cells were imaged with a fluorescence microscope. **A.** Fluorescence imaging of CAD cells either showed tip accumulation (V144F, V220I, E233D, and A255V) comparable to WT, Partial tip accumulation (S58L, A202P, R316W, and T312M) significantly less than WT, cell body retention of motors (R216P, R216H, and L249Q) or microtubule decoration (T46M, T99M, G102D, S215R, and E253K), the tip of the axon are marked by yellow arrowhead **B.** Bar graph representing peripheral accumulation (FI at the tip/ FI in cell body) for WT and mutant motors as mean ± SEM (two-tailed unpaired t-test. **** p < 0.0001, * p 0.0114, ns: non-significant).

Furthermore, CAD cell expression of microtubule-binding mutants T46M, T99M, G102D, S215R and E253K exhibited strong microtubule decoration as seen in COS-7 cells (**Fig. 3A**). Interestingly, these mutants inhibited CAD cell differentiation into the characteristic cell body and neurite instead showed round morphology (**Fig. 3A**); presumably, the binding of motors affected the microtubule dynamics critical for differentiation of neurons. The data suggests that microtubule dynamics and neuronal transport play a critical role in neuronal growth, differentiation and function. Collectively, these results indicate that most KIF1A neurodegenerative disease mutations have an adverse effect on KIF1A motor function either by affecting the motor velocity, processivity and microtubule affinity or by locking the motor on the microtubule.

### KIF1A disease mutants exhibit distinct motility properties

To ensure that the observed phenotypic distribution of disease mutants in COS-7 and CAD cells reflects their motility properties, we performed TIRF microscopy-based single-molecule motility assays. In general, disease mutations in the motor domain decreased the motor velocity and landing rate compared to WT motors (**Fig. S4B**). The truncated dimeric, WT KIF1A(393LZ) motor displayed long superprocessive motility events along the microtubule surface in the presence of ATP (**Fig. 4**) with an average velocity and run-length of 2.48 µms^-1^ and 10.26 µm, respectively (**Fig. 4; Fig. S4; Table 2**). Similarly, motility analysis of peripheral mutants displayed long processive runs along the microtubule surface comparable to that of WT motors, albeit with less frequency. Motility analysis of V144F mutant showed microtubule-based long processive motility along with an average velocity and run-length of 0.62 µms^-1^ and 4.88 µm, respectively (**Fig. 4; Fig. S4; Table 2**). The mutations V220I and E233D resulted in average velocities and run-lengths of 1.22 µms^-1^ and 7.79 µm and 0.49 µms^-1^ and 7.17 µm, respectively (**Fig. 4; Fig. S4; Table 2**). Mutation A255V, though the motors showed peripheral accumulation in the COS-7 cells. Surprisingly, in the single-molecule motility assay, the mutant showed slow back-and-forth diffusive motion along the microtubule surface (**Fig. 4**).

**Figure 4:**
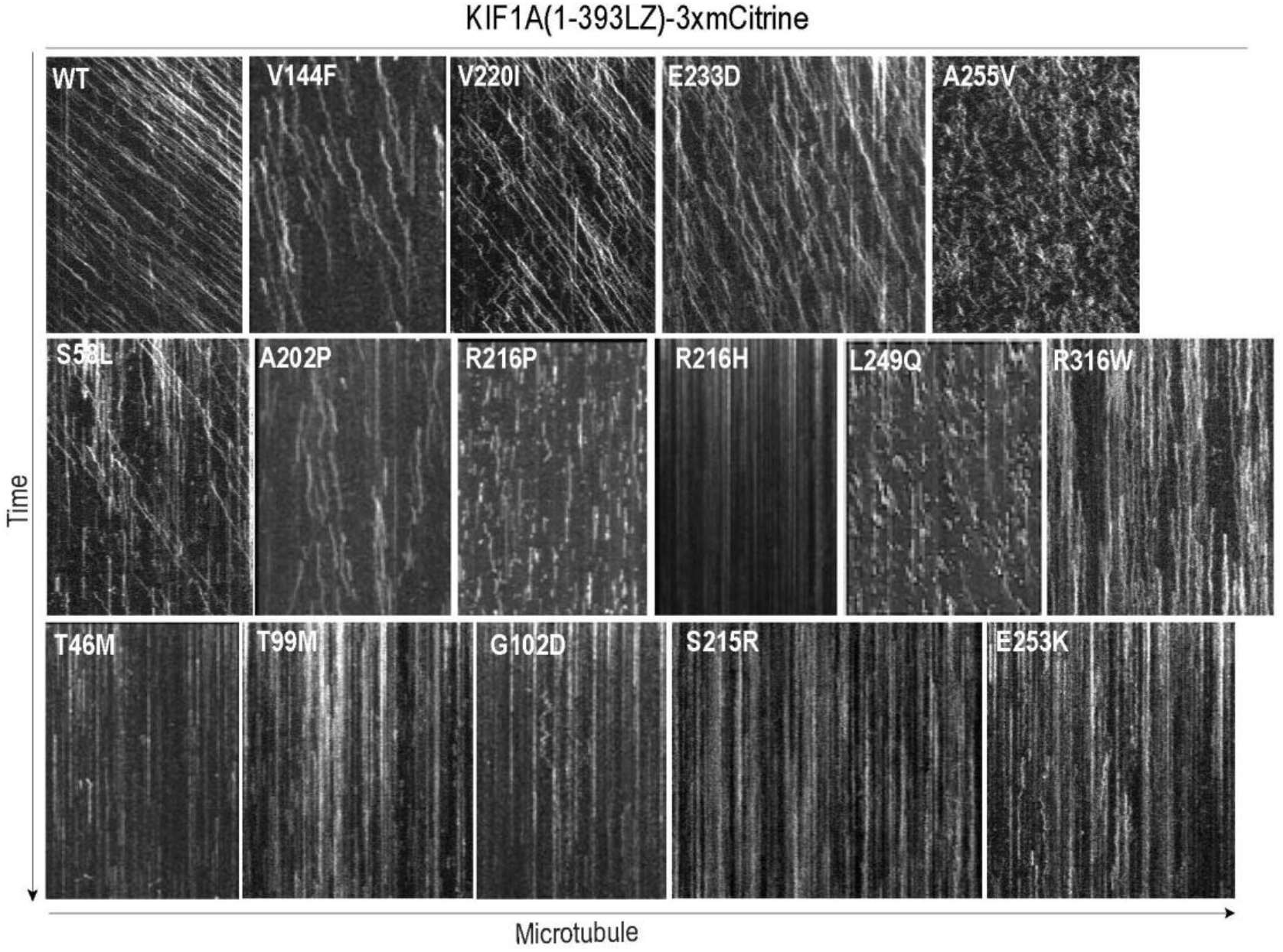
*In vitro* single molecule motility assay. COS-7 cell lysate was flown onto the microtubules adsorbed on a glass coverslip, and motility events were recorded using TIRF microscopy. Representative kymographs for each mutant are shown with microtubule length on X-axis and time on Y-axis.

**Table 2:** Motility properties of KIF1A WT and mutant motors in *in vitro* assays

| No. | KIF1A<br>(1-393LZ)-<br>3xmCitrine | SMMA | SMMA<br>velocity | SMMA<br>run length | Gliding<br>velocity |
| --- | --- | --- | --- | --- | --- |
| 1 | WT | Motile | $2.48 \pm 0.02 \mu\text{m/s}$ | $10.26 \pm 0.25 \mu\text{m}$ | $1.28 \mu\text{m/s}$ |
| 2 | V144F | Motile | $0.62 \pm 0.01 \mu\text{m/s}$ | $4.88 \pm 0.11 \mu\text{m}$ | $0.42 \mu\text{m/s}$ |
| 3 | V220I | Motile | $1.22 \pm 0.01 \mu\text{m/s}$ | $7.79 \pm 0.27 \mu\text{m}$ | $1.22 \mu\text{m/s}$ |
| 4 | E233D | Motile | $0.49 \pm 0.01 \mu\text{m/s}$ | $7.17 \pm 0.22 \mu\text{m}$ | $0.83 \mu\text{m/s}$ |
| 5 | A255V | Binds and Jiggles | | | $1.24 \mu\text{m/s}$ |
| 6 | S58L | Low processive | $1.19 \pm 0.02 \mu\text{m/s}$ | $2.11 \pm 0.14 \mu\text{m}$ | |
| 7 | A202P | Low processive | $0.51 \pm 0.01 \mu\text{m/s}$ | $1.71 \pm 0.05 \mu\text{m}$ | |
| 8 | R216P | Low processive | $0.38 \pm 0.01 \mu\text{m/s}$ | $0.59 \pm 0.04 \mu\text{m}$ | |
| 9 | R216H | Non-motile |  |  |  |
| 10 | L249Q | Non-motile |  |  |  |
| 11 | T312M | Non-motile |  |  |  |
| 12 | R316W | Binds and Jiggles | $0.43 \pm 0.01 \mu\text{m/s}$ | $0.59 \pm 0.3 \mu\text{m}$ | |
| 13 | T46M | Non-motile |  |  |  |
| 14 | T99M | Non-motile |  |  |  |
| 15 | G102D | Non-motile |  |  |  |
| 16 | S215R | Non-motile |  |  |  |
| 17 | E253K | Non-motile |  |  |  |

We next performed similar single-molecule motility assays with cytoplasmic mutants. The S58L mutant showed short processive motility along the microtubule with an average velocity of 1.19 µms^-1^ and run-length of 2.11 µm (**Fig. 4; Fig. S4; Table 2**), consistent with its cytoplasmic distribution in COS-7 assay and neurite tip accumulation in CAD cells. Similarly, A202P also exhibited slow and short processive runs along the microtubule with an average velocity of 0.51 µms^-1^ and run-length of 1.71 µm (**Fig. 4 Fig. S4; Table 2**). However, mutant R216P showed slow back-and-forth diffusive motion along the microtubule with an average velocity of 0.38 µms^-1^ and run length of 0.59 µm (**Fig. 4: Fig. S4; Table 2**). For R316W mutant, consistent with its cytoplasmic localization in COS-7 assay and partial neurite tip accumulation in CAD cells, the motor showed slow (0.43 µms^-1^) and short (0.59 µm) back-and-forth motion on the microtubule surface *in vitro* (**Fig. 4 Fig. S4; Table 2**). However, R216H, L249Q and T312M mutants completely lost microtubule-binding and motility (**Fig. 4**).

Together these results suggest that mutations that showed peripheral localization in COS-7 cells showed a minor effect on KIF1A motility properties and showed comparable velocity but lower run length than WT motors. In contrast, S58L, A202P and R316W mutants showed partial neurite tip accumulation in CAD cells, displayed low velocity and reduced run length. Finally, complementing their microtubule-bound phenotype in COS-7 and CAD cell assays, the single-molecule analysis of T46M, T99M, G102D, S215R and E253K mutants failed to generate motility events along the microtubule surface (**Fig. 4**).

### KIF1A disease mutants display variable multi-motor microtubule gliding properties

Intracellular cargo transport is a highly orchestrated event requiring the collective action of multiple motors of the same or opposite polarity coordinating their activities in ensembles. Thus, we employed a multi-motor gliding assay to provide insights into the effect of disease mutations on the collective motion of microtubules driven by multiple WT or mutant motors. In the gliding assay, microtubules propelled over a lawn of motor proteins coated onto the cover glass as described (Soppina et al., 2022a; Soppina et al., 2022c). Briefly, adsorb the flow chamber with GFP nanobodies followed by mCitrine-tagged motors to coat the motors on the surface. Rhodamine-labeled microtubules were sheared before infusing into the motility chamber, and gliding events were recorded using TIRF microscopy (**Fig. 5A**).

**Figure 5:**
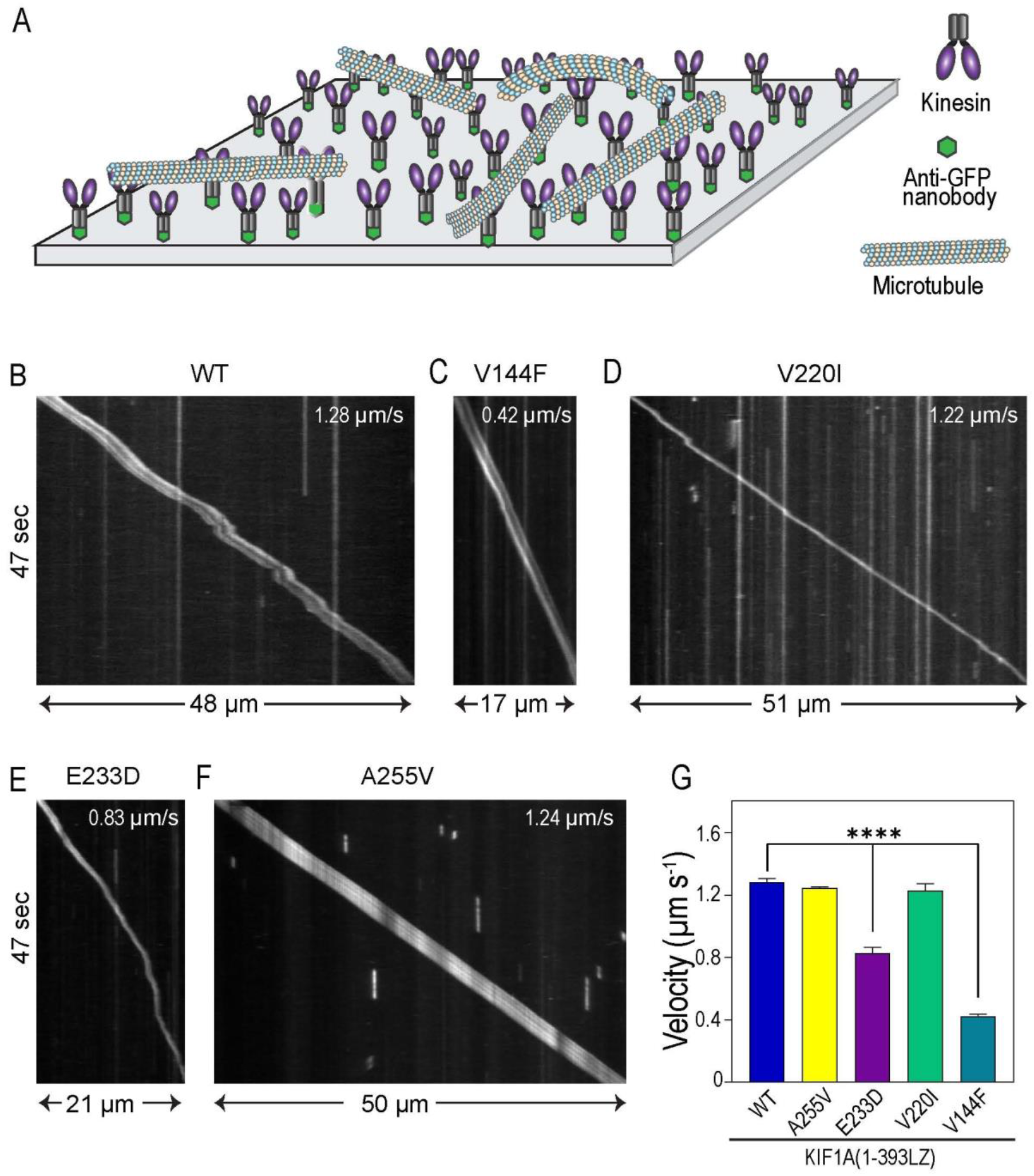
Multi-motor gliding by motile KIF1A mutants. **A.** Cartoon representation of rhodamine-labeled microtubules gliding on the mCitrine-tagged motors bound to glass surface via GFP-nano body. Representative kymograph of microtubule-gliding event for the constitutively acive **B.** WT, **C.** V220I**, D.** E233D**, E.** E233D**, F.** A255V mutant motors, **G.** Bar-graph showing gliding velocity calculated from three independent experiments as mean±SEM for WT and mutants (two-tailed unpaired t-test. **** p < 0.0001)

The R216H, L249Q, and T312M mutants completely lost their ability to interact with the microtubule, whereas T46M, T99M, G102D, S215R, and E253K showed strong microtubule-binding in single-molecule motility assays. Therefore, we did not perform microtubule gliding analysis for these mutants. The microtubule gliding assays for peripheral mutants (V144F, V220I, E233D, and A255V) showed multiple gliding events and their gliding velocities were calculated using velocity measurement tool macro in Fiji/ImageJ. The WT motor showed a mean microtubule gliding velocity of 1.28 µms^-1^ (**Fig. 5, B and G**; **Movie 1**; **Table 2**), consistent with previous studies (Soppina et al., 2022a; Soppina et al., 2022c). However, gliding velocity of V144F and E233D mutants were found to be 0.42 µms^-1^ and 0.82 µms^-1^, respectively, which is significantly lower than WT motors (**Fig. 5, C, E and G; Movie 2 and 3**; **Table 2**). These mutations considerably impacted motor coordination, essential for efficient microtubule gliding. This follows the single-molecule motility data where V144F and E233D showed lower velocity and run length than WT motors. V220I and A255V showed microtubule gliding velocities of 1.25 µms^-1^ and 1.22 µms^-1^, respectively (**Fig. 5, D, F and G: Movie 4 and 5**; **Table 2**). Though these microtubule gliding velocities were comparable to WT motors, mutants showed lower velocities in single-molecule motility assay, possibly due to the collective efforts of the mutant motors. Interestingly, gliding analysis with A255V mutant displayed frequent and significant microtubule bending and spiraling (**Movie 5**) compared to WT motors (Soppina et al., 2022a).

Together the data suggest that V144F and E233D mutations presumably disrupt motor coordination between the heads, consistent with measured lower velocities in single-molecule motility and multi-motor gliding assays. In contrast, A255V mutant that showed back-and-forth motion in single-molecule motility assay could glide the microtubules as efficiently as WT motors. On the other hand, V220I mutant did not show a considerable difference from WT in collective motor behavior. These results agree with previous microtubule gliding studies for A255V and V220I mutations (Esmaeeli Nieh et al., 2015).

### KIF1A disease mutants show altered low load cargo transport *in vivo*

Collective dynamics of multiple motors of similar or different types are essential for generating higher forces and regulating cellular cargo navigation in many biological processes. Defects in cargo transport have been strongly correlated with several neuronal diseases. Thus, we wanted to know whether disease mutants of KIF1A motors can work together in teams to overcome their motility defects and drive long-distance cargo transport in cells (Ucar and Lipowsky, 2020; Verhey et al., 2011).

To examine this, we employed a non-native peroxisome dispersion assay (Kapitein et al., 2010; Schimert et al., 2019). Peroxisomes are single membrane-bound stationary organelles with sizes ranging from 20 to 250 nm in diameter localized at the perinuclear region (Soliman et al., 2018) and require <15-pN force for their dispersion; thus, we consider them as low-load cargo (Efremov et al., 2014). This assay facilitates the analysis of multi-motor behavior in a physiological environment where motors work in teams to transport membrane-bound cargo. Thus, KIF1A WT or mutant motors are targeted to the peroxisome membrane, resulting in motion and cellular distribution of these organelles, attributed to motor’s ability to transport cargo (**Fig. 6A**).

**Figure 6:**
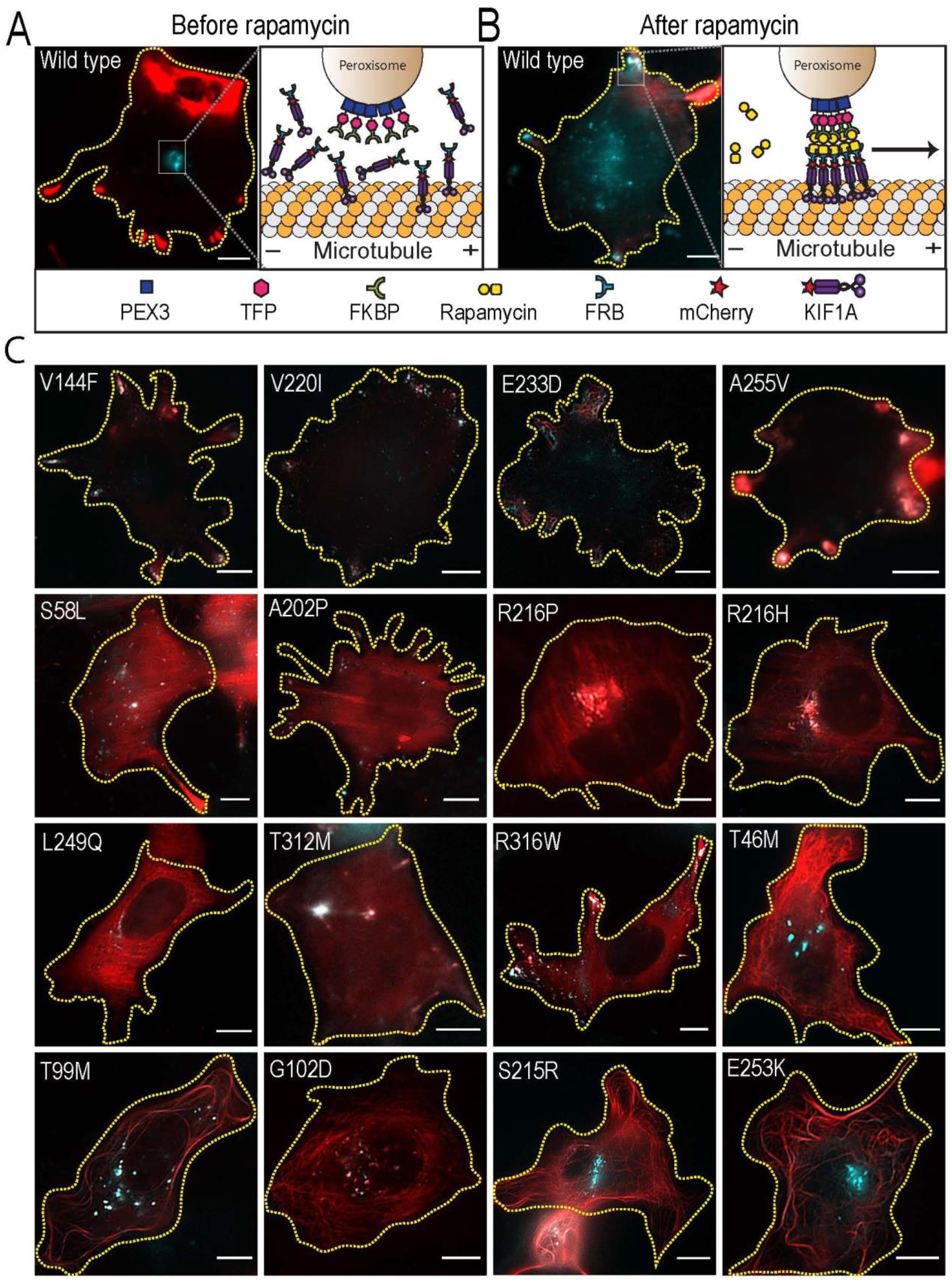
Peroxisome transport by WT and mutant KIF1A motors. Graphical representation of peroxisome tagged with PEX3-TFP-FKBP (cyan) and **A.** unloaded KIF1A-mCherry-FRB motors (red) before addition of rapamycin and **B.** peroxisome loaded with motors walking on the microtubule on addition of rapamycin, **C.** Merged image of WT and mutant motors expressed in COS-7 cells targeted to peroxisomes and imaged after 30 minutes of rapamycin treatment. Scale bar: 10µm.

For this assay, a fluorescent peroxisome targeting module (PEX3-mTFP-FKBP) was assembled by fusing a peroxisome targeting peptide with a teal-fluorescent protein (TFP) and FKBP (FK506 binding protein. Similarly, we tagged the C-terminus of WT and disease-mutant KIF1A motors with mCherry and FRB (FKBP-rapamycin binding domain). We coexpressed the tagged motors with a peroxisome-targeting module in COS-7 cells. We observed no visible motor recruitment onto the peroxisomes in the absence of rapamycin (**Fig. 6A**). Rapamycin or FKBP12 alone has a very low binding affinity for FRB, whereas FKBP-rapamycin complex has approximately 2000-fold higher affinity for FRB (Banaszynski et al., 2005). Thus, adding rapamycin triggers rapid FRB-FKBP heterodimer complex formation, resulting in motors loading onto the peroxisome surface (**Fig. 6A**). To determine the effect of KIF1A disease mutations on *in vivo* cargo transport, we performed a quantitative analysis of the motor/cargo dispersion before at 0 min and after 30 min of rapamycin treatment by fluorescence microscopy (**Fig. S5**).

For WT motor, upon rapamycin treatment, 27.3% of cells exhibited partial peroxisome dispersion remaining 72.7% showed peripheral localization (**Fig. 6A; Fig. S5, A and B; Fig. S6**). For V144F, consistent with decreased velocity, processivity and microtubule landing in the motility assays (**Fig. 6C**), only 32.2% of cells showed robust peroxisome dispersion to the periphery and 67.8 % of cells showed partial dispersion (**Fig. 6C; Fig. S5, A and B**) compared to WT motors. As V220I mutant exhibited similar motility properties to WT motors, the mutant showed peripheral peroxisome distribution in 65.2% of cells and 34.8% with partial distribution (**Fig. S5, A and B; Fig. S6**), comparable to WT motors. Similarly, E233D mutation showed a minor effect on motor processivity and a drastic effect on motor velocity and microtubule affinity; therefore, these mutants exhibited 45.9% of cells with peripheral localization and the rest, 54.1% cells showed partial dispersion of peroxisomes (**Fig. 6C; Fig. S5, A and B; Fig. S6**). Furthermore, although A255V mutant displayed diffusive motility in single-molecule assays, motors dispersed the peroxisomes to the cell periphery in 49.3% of cells and partial dispersion 51.7% of cells (**Fig. 6C; Fig. S5, A and B; Fig. S6**). Interestingly, these results complement with measured smooth and uniform microtubule gliding with an average velocity comparable to WT motors, suggesting that mutant motors collectively work in teams for efficient cargo transport.

Compared to WT motors, recruitment of cytoplasmic mutant S58L partially dispersed peroxisomes in 88.9% of cells and 11.1% showed perinuclear distribution, consistent with its decreased velocity, microtubule affinity and processivity in single-molecule motility assays and partial tip accumulation in CAD cells. Similarly, though the cytoplasmic mutant A202P showed decreased velocity, processivity and microtubule affinity in single-molecule assays, the mutant showed 28.6% peripheral, 71.4% partial, and 12.9% perinuclear distribution of peroxisomes (**Fig. 6C; Fig. S5, A and B; Fig. S7**). L249Q, a cytoplasmic mutant failed to generate processive motility in single-molecule and CAD cells assays, also failed to disperse peroxisomes in 50% of cells and remaining 50% cells showed perinuclear distribution (**Fig. 6C; Fig. S5, A and B; Fig. S7**). T312M, a cytoplasmic mutant that failed to generate microtubule-based processive motility in single-molecule motility assays also was unable to disperse peroxisomes in 22.4% of cells but showed perinuclear distribution in 64.7% of cells and partial dispersion in 12.9% of cells (**Fig. 6C: Fig. S5, A and B; Fig. S7**). Similarly, R316W, a cytoplasmic mutant that exhibited diffusive motility at the single-molecule level and partial tip accumulation in CAD cells, dispersed the peroxisomes to cell periphery in 20.6% of cells, partial distribution in 55.1% and perinuclear localization in 24.3% of cells (**Fig. 6C; Fig. S5, A and B; Fig. S7**). Similar targeting of T46M, T99M, G102D, S215R and E253K mutants strongly decorated microtubules in COS-7, CAD and single-molecule assays, to peroxisomes failed to induce peroxisome dispersion (**Fig. 6C and Fig. S8**). Instead, these cargo-bound mutants decorated the microtubules, presumably due to their impaired catalytic cycle (**Fig. S8**).

Together, the data suggest that although V144F, V220I, E233D and A255V mutations had a mild effect on motor superprocessivity and significantly reduced microtubule affinity, mutant motors collectively work in teams and efficiently transport cellular cargoes. Similarly, S58L, A202P, L249Q, T312M, and R316W mutations severely affected KIF1A motility and microtubule binding and resulted in impaired cellular cargo transport. Furthermore, mutations R216H and R216P completely abolished motor-microtubule interaction and consequently failed to transport cellular cargoes.

### KIF1A disease mutants have a drastic effect on force generation *in vivo*

The kinesin motor domain executes microtubule binding, processive motility and force generation. Despite a high level of sequence and structural conservation, we have shown previously that kinesin-3 family motors are inherently fast and supreprocessive with a strong affinity for the microtubule (Scarabelli and Grant, 2013; Soppina et al., 2022b; Soppina et al., 2022d; Soppina et al., 2014; Soppina and Verhey, 2014). However, the ability of kinesin-3 motors to generate force under opposite loads is poorly studied. Much of the current understanding of how kinesin motors generate force mostly comes from biophysical analysis of kinesin-1 motors. Studies have shown that slight variations in subdomains of the motor head determine family-specific properties. In fact, recent force measurement studies on mammalian KIF1A exert a maximal force of ∼3pN with frequent detachment from the microtubule. However, the family-specific K-loop insert in the loop12 of the motor domain enables swift reattachment to the microtubule and resumes force generation (Soppina and Verhey, 2014). Nevertheless, the effect of disease mutations in the KIF1A motor domain on its ability to generate force under the opposite load has never been tested. Thus, we examined the impact of disease mutation on KIF1A force generation.

To test this, we employed a Golgi dispersion assay; WT or mutant motors were targeted to the Golgi by fusing Golgi targeting sequence HsGMAP210 (amino acids 1757-1838, NP_004230) to the C-terminal region of the motor. Golgi is a non-native heavy load cargo that requires 100 to 200 pN of opposite forces to disrupt its tight perinuclear localization **(Fig. 7A)**. The WT KIF1A(1-393LZ)-mCit-GTS motors showed complete dispersal of Golgi towards the periphery **(Fig. 7B)**. However, targeting motors carrying disease mutations showed partial to no dispersion of Golgi from its tight perinuclear localization. The Golgi targeting of peripheral mutants, V144F, V220I, E233D and A255V, exhibited distinct Golgi scattering in cells (**Fig. 7C; Fig. S9, A and B; Fig. S10**) compared to WT motors **(Fig. S9, and B)**. For V144 mutant, the Golgi loading resulted in 8.6%, 72.8% and 18.5% of cells with a peripheral, partial and perinuclear scattering of Golgi (**Fig. 7C; Fig. S9, A and B; Fig. S10**), respectively. These results agree with the partial dispersion of peroxisomes and measured decrease in the motor velocity, processivity and microtubule affinity in single-molecule assays. Similarly, recruitment of E233D or V220I mutants resulted in a partial distribution of Golgi in 76.3% and 84% of cells, respectively and peripheral accumulation in 23.7% and 15.7% of cells, respectively (**Fig. 7C; Fig. S9, A and B; Fig. S10**). Although E233D mutant exhibited slower velocity than V220I, the mutant showed efficient Golgi scattering; presumably, V220 is a conserved residue that plays an important role in force generation under high load. Interestingly, loading A255V mutant with diffusive motility resulted in a partial distribution of Golgi in 60% of cells and 40% of cells with peripheral Golgi scattering. Collectively, the data suggest that although peripheral mutations have a minor effect on motor’s processivity, they significantly impact the ability to generate higher force compared to WT motors. Though A255V mutant showed slow diffusive motion at single-molecule assays, the mutant effectively scattered the Golgi, suggesting that this conserved residue in the Switch-II is critical for generating force under high load.

**Figure 7:**
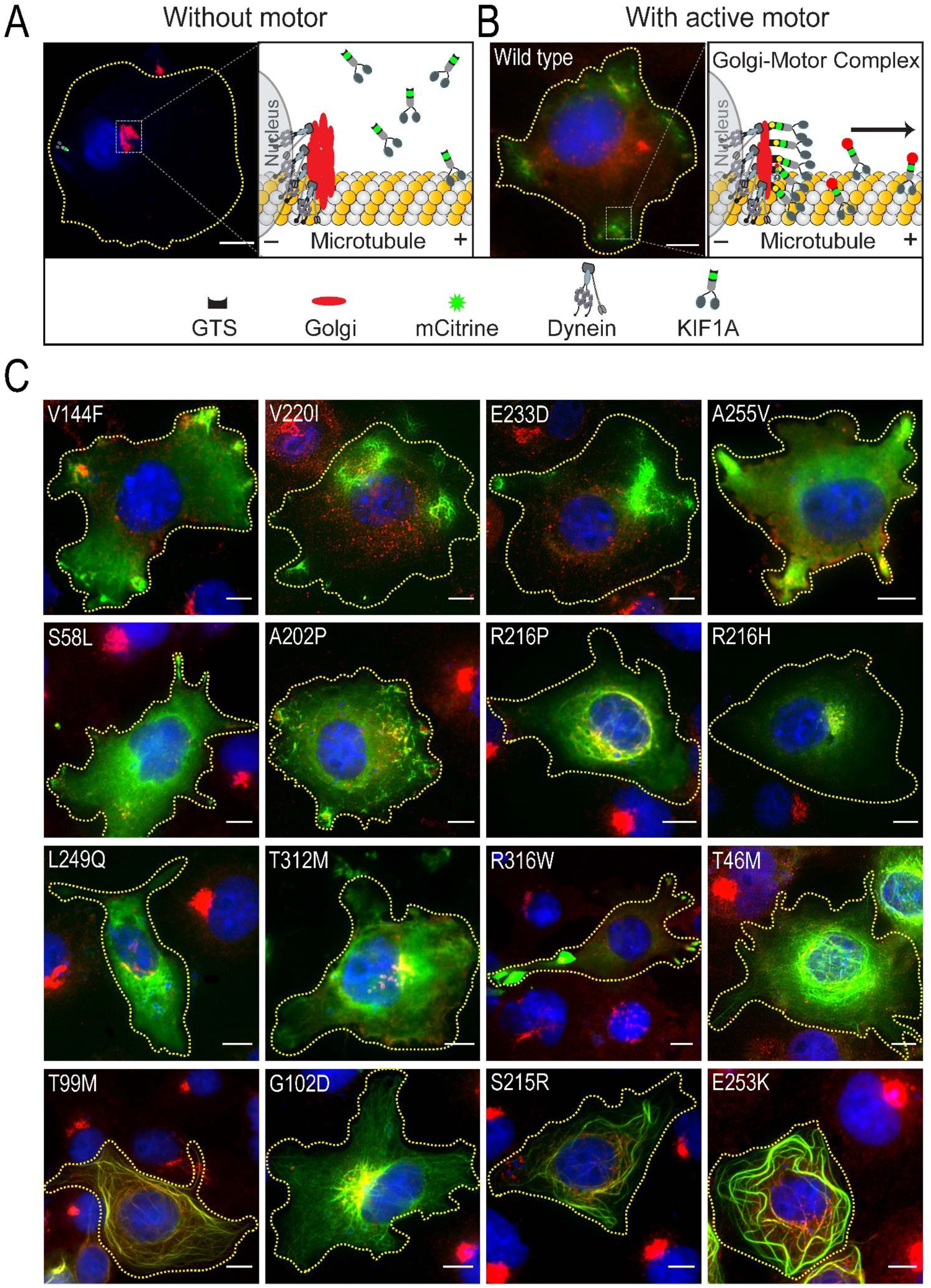
Transport of heavy-load cargo (Golgi) by WT and mutant KIF1A motors. Cartoon representation of Golgi **A.** without any motor expression and with **B.** WT motor. **C.** Representative TIRF image showing the KIF1A WT and mutant motors (green) tagged with GTS sequence expressed in COS-7 cells. Blue is nuclear staining with DAPI dye, and the red display Golgi stained with an anti-giantin antibody. Scale bar: 10µm.

The Golgi recruitment of cytoplasmic mutant S58L showed perinuclear scattering in 72% of cells, while 28% of cells showed intact and tight Golgi (**Fig. 7C; Fig. S9, A and B; Fig. S11**), consistent with its moderate processivity and partial peroxisome dispersion. Next, targeting A202P, a low processive cytoplasmic mutant, partially scattered the Golgi in 94% of cells and only 6% of cells showed complete Golgi scattering (**Fig. 7C; Fig. S9, A and B; Fig. S11**). Similar analysis of cytoplasmic mutant R216H failed to disperse the Golgi in 85% of cells and 15% showed slight perinuclear scattering, consistent with its single-molecule data where the mutant failed to interact with microtubules (**Fig. 7C; Fig. S9, A and B; Fig. S11**). Furthermore, the loading of R216P, a cytoplasmic mutant that has microtubule binding but failed to generate processive motility *in vitro*, caused 16 % peripheral and 66% perinuclear Golgi scattering and the remaining 18% of cells showed intact Golgi (**Fig. 7C; Fig. S9, A and B; Fig. S11**). Similar recruitment L249Q and T312M failed to scatter the Golgi in 93% and 78% of cells (**Fig. 7C; Fig. S9, A and B; Fig. S11**), respectively, consistent with their inability to interact with microtubules in the single-molecule assays. The Golgi targeting of cytoplasmic mutant R316W that showed back-and-forth diffusive motion along the microtubule scattered the Golgi partially in 87% of cells (**Fig. 7C; Fig. S9, A and B; Fig. S11**). As T46M, T99M, G102D, S215R and E253K showed strong microtubule binding, loading of these mutants resulted in strong microtubule decoration of Golgi fragments at the perinuclear region (**Fig. 7C; Fig. S9, A and B; Fig. S12)**.

Together, the data indicate that peripheral mutations moderately affected motor velocity and force generation compared to wild-type motors. While cytoplasmic mutations strongly affected motor velocity, processivity, and their ability to generate force. Furthermore, microtubule-binding mutations not only abolished motility properties but also abolished their ability to generate force. Determining the exact force generated by each mutant motor requires optical trap-based measurements at the single-molecule level.

### Disease mutation in full-length KIF1A hamper native cargo transport

KIF1A is known to transport a variety of cellular and neuronal cargoes, such as synaptic vesicle precursor, neurofilament, and dense-core vesicles critical for neuronal development and signaling. We hypothesized that the decreased velocity and landing rate of the mutant motors would hinder the efficiency of cargo transport in cells. However, it remained possible that multiple motors on the cargo could work together in teams and negate the effect of disease mutation. To test this, we expressed a full-length version of WT or V144F mutant motor in COS-7 cells and analyzed their cargo transport properties in cells. The cells expressing full-length KIF1A WT or mutant motors showed vesicular structures with robust microtubule-based bidirectional motion, while microtubule depolymerization resulted in a complete loss of vesicle motion. The WT KIF1A positive vesicles traveled at a velocity of 2.07±0.04 µms^-1^ **(Fig. 8A and Movie 6)**. In contrast, the mutant KIF1A (V144F) transported the cargoes at a speed of 0.97±0.02 µms^-1^, which is significantly lower than WT motors **(Fig. 8B and Movie 7)**. The *in vivo* cargo velocities of WT and mutant motors significantly correlate with their velocities measured at single-molecule levels *in vitro*. As the mutant KIF1A motor walks two times slower than the WT motor, it would take longer to deliver the cargo to its destination (**Fig. 8C**). Together; the data suggest that though V144F mutation did not significantly affect the motor processivity, it drastically reduced the velocity and microtubule affinity, presumably the cause of neurodegeneration. This shows that our findings with constitutively active KIF1A motors can be extrapolated to full-length KIF1A mutant motors.

**Figure 8:**
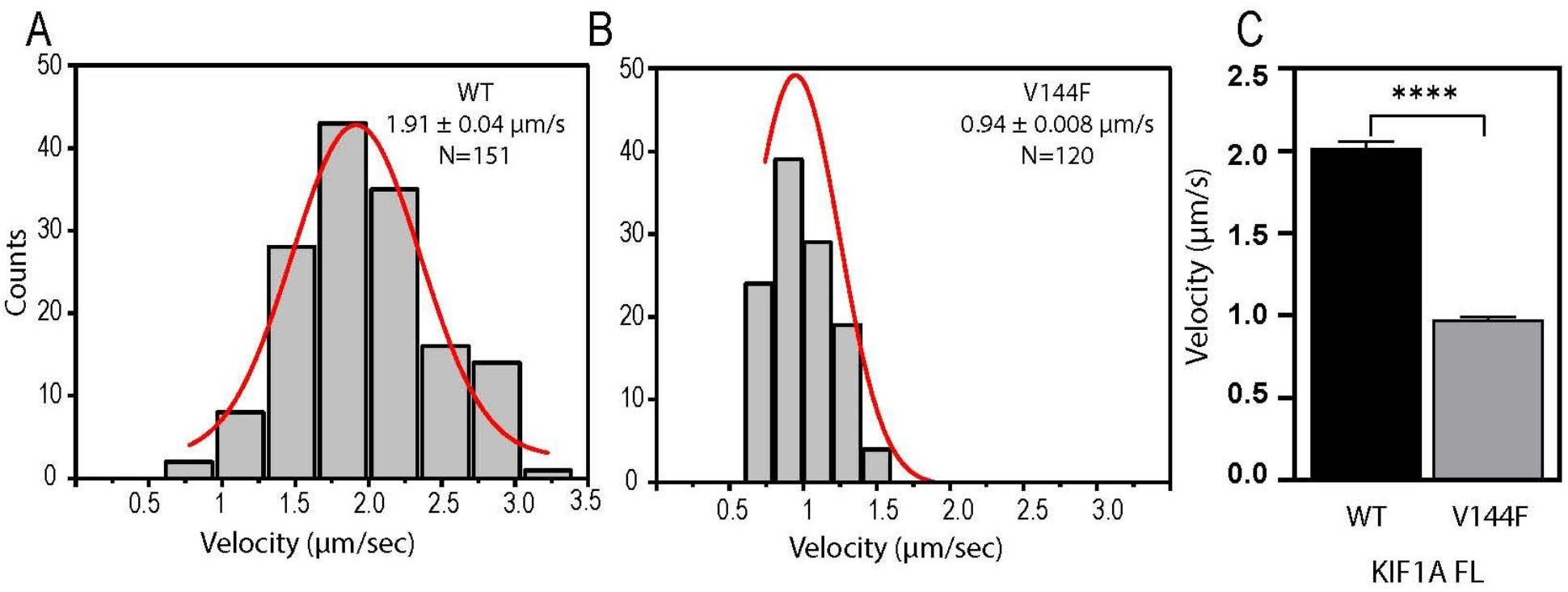
In-vivo transport of native cargo by full-length KIF1A WT and V144F mutant motors. Histogram displaying the range of velocities fit to Gaussian distribution for vesicular transport by **A.** full-length WT motor and **B.** V144F mutant. The velocity and number of events (N) plotted are displayed in the right corner of the histogram. **C.** Bar graph showing velocities for WT and V144F mutant motor. Data was tested by unpaired t-test (two-tailed, **** p < 0.0001)

## Discussion

### A novel cell-based assay to study the effect of mutations on KIF1A motility properties

We have demonstrated that kinesin-3 motors exhibit peripheral accumulation in COS-7 cells due to their superprocessive motility and high landing rates. In contrast, mutants with short processive motility are distributed uniformly in the cytoplasm. The assay can be used to investigate the mechanism of kinesin-3 superprocessivity by generating a large number of mutants using random mutagenesis. Furthermore, it would be a great tool for high throughput screening of potential small molecules that modulate the activity of kinesin-3 motors. The only caveat is the assay is not sensitive to determine the change in the motor velocity due to disease mutation. Here we employed this assay to perform a comprehensive characterization of 16 KIF1A neurodegenerative disease mutations, including 12 heterozygous (V144F, S58L, A202P, R216H, R216P, L249Q, R316W, T99M, G102D, S215R, E253K and T312M), 2 homozygous (A255V and R350G) and 3 polymorphic (T46M, V220I and E233D).

### Motility properties of peripheral mutants correlate with mild disease symptoms

The mutations of V144F, V220I, E233D and A255V showed peripheral accumulation of motors in the COS-7 cells, consistent with the *in vitro* single-molecule motility analysis showing a little-to-no effect on KIF1A cargo transport, force generation and motility properties (**Fig. 4; Fig. S4; Table 1**). Intriguingly, these mutations significantly decreased the motor velocity by 2-3 fold (**Fig. S4 and Fig. 8C**). The residue V144 is highly conserved on the β4-strand plays a crucial role in ATP binding and hydrolysis during processive motility. In amino acids, the structure of the side chain determines the steric interaction, hydrophobic volume, and buried surface area, which are essential to the hydrophobic core stability (Munson et al., 1996). Thus, the mutation V144F likely destabilizes the α3 helix of switch I region, critical for ATP binding or γ -phosphate release of the motor domain, due to its bulky side chain and explains the substantial decrease in motor velocity and processivity compared to WT motor (Percy and McMullin, 2005). Similarly, V220I mutation on the β6 strand slightly decreased KIF1A velocity with little to no effect on motor processivity and microtubule affinity. The β6 connects the central β sheet to switch-I and communicates the conformational changes to the switch-II region due to ATP hydrolysis. The mutation from valine to isoleucine slightly increased hydrophobic volume due to its longer side chain, resulting in a small effect of the V220I mutation on KIF1A velocity (Behmard et al., 2012). The mutation, E233D, in the loop10 (L10) of KIF1A motor domain also did not alter the motor’s processivity but drastically decreased the motor velocity compared to the WT. There could be two possibilities; first, the mutation did not alter the overall charge or the hydrophobicity in the loop. Second, swapping loop10 (L10) in KIF7 with that of kinesin-1 did not alter its microtubule-binding affinity, suggesting L10 may have no major role in microtubule binding and motility (Jiang et al., 2019).

Akin to the wild type, the A255V mutant showed peripheral localization in COS-7 and tip accumulation in CAD cells. However, in single-molecule assays, unlike previously reported fast and hyperactive motility (Chiba et al., 2019; Guedes-Dias et al., 2019), mutant motors showed slow diffusive motion along the microtubule surface. Valine is a strong hydrophobic residue primarily present in the protein interior to determine the protein structure and stability (Fitzsimmons et al., 2016). Its presence on the protein surface contributes to defective folding, presumably why motors exhibited back-and-forth motion along the microtubule surface. A255V mutation in the L11 probably disrupted motor-microtubule interaction in ATP state, but motors utilized a positively charged K-loop in the L12 to tether and diffuse along the microtubule through an electrostatic interaction with the negatively-charged microtubule surface (Okada et al., 2003; Okada and Hirokawa, 1999; Okada and Hirokawa, 2000; Soppina and Verhey, 2014). In the ATP –bound state, KIF1A extends its L11 down and interacts with the microtubule to generate processive motility (Nitta et al., 2004). Upon ATP-hydrolysis, ADP-bound KIF1A retracts L11, extends down its positively charged L12 and interacts with the microtubule, resulting in diffusive motion. Remarkably, in the multi-motor gliding assay, the mutant showed smooth microtubule gliding with significant microtubule bending is interesting. We have recently demonstrated that the K-loop, the family-specific insertion in the L12 of kinesin-3 motors, induces microtubule bending (Soppina et al., 2022a). However, A255V mutants inducing microtubule-bending need further detailed investigation to understand the underlying cause at the mechanistic level.

In conclusion, the data suggest that peripheral mutants, V144F, V220I, E233D and A255V, exhibit mild effects on KIF1A motility properties compared to wild-type, likely the reason patients suffering from neurodegenerative diseases due to these mutations in the motor domain display mild clinical phenotypes (Gabrych et al., 2019; Klebe et al., 2012; Lee et al., 2015).

### Motility properties of cytoplasmic and microtubule-binding mutants correlate with severe disease phenotype

Next, among the mutations that showed cytoplasmic localization in COS-7 cells, mutations of S58L, A202P, R216H and R216P drastically affected their microtubule affinity and motility properties. However, mutations of L249Q, R316W, and R350G showed microtubule binding but failed to generate motility along the microtubule. S58L, a dominant, heterozygous mutation in the β2, likely interacts with conserved tyrosine (Y67) residue in the loop3 (L3), thought to be important for ATP binding. As serine (S58) residue is highly conserved in the kinesin-3 motors, mutating it to a hydrophilic residue disrupts its interaction with Y67, resulting in a remarkable decrease in the microtubule affinity, motor velocity and processivity. Similarly, mutation A202P is at switch-I junction of α3 and loop9 (L9), presumably interferes with the β-to-α conformational transition of loop9 and hinders phosphate release (Nitta et al., 2004), resulting in a slow processive motor with low microtubule affinity.

The R216P and R216H mutants loosely tethered the microtubule but failed to generate processive motility hinting that the mutation abolished the formation of a salt bridge between R216 (L9) and E253 (L11) residues, critical for phosphate release after ATP hydrolysis and generated processive motility (Nitta et al., 2004). In contrast, mutations of T312M and R350G located in α5 and α6 regions completely abolished microtubule association. Threonine (T312) is located in α5 region important for stabilizing the microtubule-binding regions loop11 and loop12. The T312M mutation likely abolishes its interaction with loops and affects microtubule binding. For R350G mutation, we could not perform microtubule-based motility assays because the mutant barely expressed in mammalian cells and it took almost 48h to see the protein expression.

T46M, T99M, G102D, S215R and E253K disease mutants showed strong microtubule-binding but failed to exhibit processive motility along the microtubule. These mutations are located in the ATP-binding pocket of KIF1A and are critical to establishing interactions with ATP. Therefore, any alterations to these residues presumably disrupt ATP binding or prevent ATP hydrolysis and the motor gets locked on the microtubule in a no-nucleotide state. These results are consistent with previous structural prediction using protein modeling, suggesting that the residues of T46M, T99M and S215R mutations abolish the interactions of KIF1A with the γ-phosphate of ATP. However, G102D mutation in the conserved P-loop (Snider and Houry, 2008) is predicted to contribute to a large structural change in the loop, thus disrupting its interaction with the γ-phosphate of ATP. As a result, these mutants bind strongly to the microtubules, stabilize the microtubules and abrogate neurite outgrowth and axonal extension. Previous studies have shown that microtubule dynamics play a key role in neuritogenesis in CAD cells and the development of neuronal processes (Li et al., 2006; Lu et al., 2004). Studies have also shown that treatment of the growth cones with vinblastine precludes microtubule dynamics and stability of the growth cone (Tanaka et al., 1995). More recent work has demonstrated that cytoskeletal elements play a crucial role in establishing the shape of neuronal and non-neuronal cells. Particularly dynamic properties of microtubule and actin filaments contribute significantly to establishing neuronal polarity (Blanquie and Bradke, 2018). Together, we show that cytoplasmic mutations considerably affected the motility properties by decreasing the velocity or processivity or decreasing/abolishing microtubule affinity, whereas microtubule-binding mutations completely abolished the motility properties by preventing nucleotide binding or release or hydrolysis by the motor domain. Notably, these motility defects directly correlate with severe neurodegenerative disease phenotypes observed in patients carrying the above mutations in the KIF1A motor domain. (Esmaeeli Nieh et al., 2015; Lee et al., 2015; Okamoto et al., 2014).

In conclusion, KIF1A-associated neurological disorders are a rare but severe form of the disease caused by a mutation in the KIF1A motor transporting precursors of synaptic vesicles in axons. The symptoms vary between the patients, even if they carry the same mutation. Our findings suggest that the microtubule-bound mutants strongly associate with the microtubules and are most inefficient in low and high-load cargo dispersion. It explains the severe disease phenotype in patients with microtubule-bound mutants of KIF1A. Similarly, the results of cytoplasmic and peripheral mutants nicely correlate with observed moderate to mild disease phenotypes.

## Methodology

### Plasmids and cloning

The constitutively active dimeric kinesin-3 construct KIF1A(1-393)-LZ-3xmCitrine has been described previously (Soppina et al., 2014). All the point mutations were generated by QuickChange site-directed mutagenesis using Turbo-Pfu polymerase (Stratagene). These mutations were subcloned into the 2xmCitrine-N1 vector with Golgi-targeting sequence (GTS) of HsGMAP210 (amino acids 1757-1838).

### Cell culture, transfection, and imaging

COS-7 (monkey kidney fibroblast, ATCC) cells were grown in Dulbecco’s modified Eagle’s medium (DMEM) containing 10% (vol/vol) fetal bovine serum (FBS; Invitrogen, India) at 37 °C with 5% (vol/vol) CO2. CAD, a mutant of a catecholaminergic cell line derived from a mouse brain tumor, was maintained in F12/DMEM with 10% FBS and 2 mM L-glutamine at 37°C and 5% CO2. These cells differentiate in serum-starved conditions. Cells were seeded at a cell density of 1 × 10^5^ in 35mm plates for microscopic imaging of fixed and live samples. The cells were transfected 24 hours post-seeding with Turbofect reagent (Thermo Fisher Scientific) according to the manufacturer’s protocol.

Image acquisition was performed with oil immersion 60X, 1.49 numerical aperture (NA) objective on an inverted Nikon Eclipse Ti2-E microscope equipped with an Andor iXon Ultra 897 electron-multiplying charge-coupled device (EMCCD) camera. A laser with 488 nm wavelength was used for exciting mCitrine fluorescent protein, a 561nm laser was used to excite mCherry fluorescent protein and a 401 nm laser was used for a teal fluorescent protein.

### COS-7 and CAD assay

COS-7 cells were seeded at a density of 1 × 10^5^ in 35mm plates and transfected with 1.0 μg KIF1A wild-type (WT) and mutant motors having C-terminal 3xmCitrine tag. CAD cells were transferred to reduced serum media prior to transfection for them to differentiate and form long axon-like structures. Both COS-7 and CAD cells were fixed approximately 12-14 hours after transfection by incubating in 4% (vol/vol) paraformaldehyde for 10 minutes and quenched by incubating in 50 mM ammonium chloride for 5 minutes. The cover glass was washed thrice with 1X PBS to remove traces of ammonium chloride and mounted on to clean the glass slide in Prolong Gold antifade reagent (Invitrogen, India). It contained 4’, 6-diamidino-2-phenylindole (DAPI) at a final concentration of 10.9 μM for staining the nuclei, and images were acquired the following day using Total Internal Reflection Fluorescence (TIRF) microscopy.

COS-7 cellular phenotype was observed in at least 50 cells for each mutant. For CAD cells, fluorescence intensity was measured using ImageJ at the cell body and tip for 30-60 cells to calculate the peripheral accumulation. The ratio of fluorescence intensity at the tip to cell body was measured peripheral accumulation. It was plotted for each mutant and compared to WT using a t-test.

### Golgi dispersion assay

We used Golgi as a heavy-load cargo to check the effect of point mutations in the motor domain on the transport by the KIF1A motor. First, we tagged the C-terminal end of the mutant motors with the Golgi-targeting sequence from the HsGMAP210 fused to the 2xmCitrine tag. COS-7 cells were transfected with these mutants, and 12-14 hours after transfection, the cells were processed for immunofluorescence. Fixing was performed as explained previously and cells were permeabilized with 0.2% Triton X-100 in PBS and blocked with 0.2% fish skin gelatin (FSG) in PBS. The Golgi apparatus was labeled with polyclonal cis-Golgi marker giantin (Poly19243, 1:2000) by incubation for 1 hour, followed by three washes, each with 0.2 % FSG. The cells were treated with secondary antibody for 1 hour and washed thrice with 0.2 % FSG and PBS. Nuclei were stained with 4’, 6-diamidino-2-phenylindole (final concentration 10.9 μM) added in ProlongGold (Invitrogen) used for mounting the coverslips. Imaging was performed after 12 hours using TIRF microscopy.

Golgi dispersion was estimated by classifying it into four categories based on the distribution of Golgi inside the cell in the presence of the mutant motor: i) tight (no dispersion, intact Golgi), (ii) perinuclear (dispersion limited to perinuclear region), (iii) partial (moderate dispersion, mostly distributed throughout cytoplasm), and (iv) complete (Golgi fragments transported to the cell periphery). These categories are represented in the schematic format in **Fig. S9A** . Values were plotted as a percentage of total transfected cells for WT and each mutant except the mutants binding the MT.

### Peroxisome scattering assay

For targeting the KIF1A motor onto the peroxisome membrane, we used a rapalog inducible FKBP/FRB heterodimerization system. Fluorescent peroxisome targeting cassette (PEX3-mTFP-FKBP) was prepared by ligating peroxisome targeting peptide with a teal-fluorescent protein (TFP) and FKBP, a FK506 binding protein. Similarly, we tagged the C terminus of wild-type and mutant KIF1A motors with mCherry and an FKBP-rapamycin binding (FRB) domain. COS-7 cells were seeded on glass-bottom dishes and cotransfected with PEX3-mTFP-FKBP and KIF1A(1-393LZ)-mcherry-FRB. Heterodimerization was induced twelve hours post-transfection by treating the cells with 1 μM rapalog. Live-cell imaging was performed at 0 and 30 min after the rapalog treatment using TIRF microscopy. For live cell imaging, cells were maintained in a live-cell chamber at 37°C with 5% CO_2_.

Peroxisome scattering was quantified by dividing the cells into the following categories: i) tight, (ii) perinuclear, (iii) partial, and (iv) complete, as described earlier. The percentage of total transfected cells was plotted for wild-type as well as mutant motors.

### Single-molecule motility assay

Single-molecule motility assay was performed using COS-7 lysate obtained by transfecting the cells with KIF1A (1-393LZ)-3xmCitrine WT and mutant motors (Soppina et al., 2022a; Soppina et al., 2014; Soppina and Verhey, 2014). Microtubules were polymerized in BRB80 [80 mM Pipes/KOH (pH 6.8), 1 mM MgCl_2_, 1 mM EGTA] with 1 mM GTP, 37°C, for 30 min using 2.5-3 mg/ml goat brain tubulin. MTs were stabilized by adding P12 buffer [12 mM Pipes/KOH (pH 6.8), 1 mM MgCl_2_, 1 mM EGTA] containing 1 μM taxol. A flow cell chamber was prepared by creating a narrow passage between two double-sided tapes stuck on the glass slide and a coverslip on the top. The MTs were diluted in P12 buffer containing taxol, flown through the chamber using a cut-tip, and allowed to adhere for 30 mins. The coverslip surface was blocked with 35-40µl of P12-BSA (15 mg/ml) buffer (with taxol) for 30 mins. Finally, motility mix containing COS-7 lysate, 15 μl of P12, 2 mM ATP, 1 mM dithiothreitol (DTT), 1 mM MgCl_2_, glucose oxidase (0.2 mg/ml), catalase (0.08 mg/ml), and 10 μM glucose was flown through the chamber. The flow chamber was sealed with wax and immediately imaged under the microscope.

Motility events were captured with oil immersion 100X 1.49 NA objective in a TIRF microscope (Nikon Ti2-E, motorized automated) and images were recorded with Andor iXon Ultra 897 EMCCD camera. Imaging was performed using a 488-nm laser at 100-ms exposure time with no delay. Fluorescent mCitrine-tagged motors moving along the length of the MT were tracked frame by frame manually to analyze the single-molecule motility properties of individual motors using a custom-written plugin for ImageJ (nih.gov). A histogram was generated by plotting the number of events for velocity and run length and fitting a Gaussian distribution for WT and mutant motors. Intermittent pauses and retraction in motility events were not considered during the measurement of velocity and run length.

### Microtubule gliding assay

The MTs were polymerized using unlabeled and rhodamine-labeled tubulin, as described in the previous section, and they were sheared into smaller fragments by pipetting up and down. GFP nanobody was flown through the flow cell and incubated for 30 minutes, followed by blocking with P12-casein buffer. The lysate-containing motors was passed through the flow cell and allowed to bind to the nanobody for 30 minutes. Finally, the motility mixture, as described earlier, along with labeled MT fragments, was passed through the flow cell. MT motility events were captured by a 561nm laser at 100ms exposure time without delay. MT gliding velocities were calculated using a macro (velocity measurement tool) in Fiji/ImageJ.

### Intracellular native cargo transport

The full-length WT and V144F mutants were expressed in COS-7 cells, and vesicles labeled with these mutants were recorded without delay using 488 nm laser in a Ti2-E microscope. These vesicles were tracked frame by frame using custom written ImageJ plugin (Soppina et al., 2022a; Soppina et al., 2014; Soppina and Verhey, 2014), and histograms were made from the number of counts for velocities of both WT and mutant motor.

## Acknowledgments

V.S. thanks Kristen J. Verhey and Roop Mallik for their unconditional support throughout the study. The authors thank all the lab members for their significant input, valuable comments, suggestions and discussions on the manuscript. V.S. acknowledges funding through DBT (Grant No.: BT/PR15214/BRB/10/1449/2015 and BT/RLF/Re-entry/45/2015) and DST-SERB (Grant No.: ECR/2016/000913). P.S. acknowledges funding from DST (Grant No.: SR/WOS-A/LS-73/2017). D.J.S acknowledges fellowship from IIT Gandhinagar.

## Author contributions

D.J.S., P.S. and V.S. designed research; D.J.S., P.S., and V.S. performed experiments; D.J.S., P.S., and V.S. analyzed data; and D.J.S., P.S. and V.S. wrote the paper with constant feedback and comments from other lab members.

## Competing interests

The authors declare no competing financial interests.

## Data and materials availability

All data are available in the main text or the supplementary materials.

## Notes

### Competing Interest Statement

The authors have declared no competing interest.

