## Supplemental file for "KIF1A neurodegenerative disease mutations modulate motor motility and force generation"

Figure S1

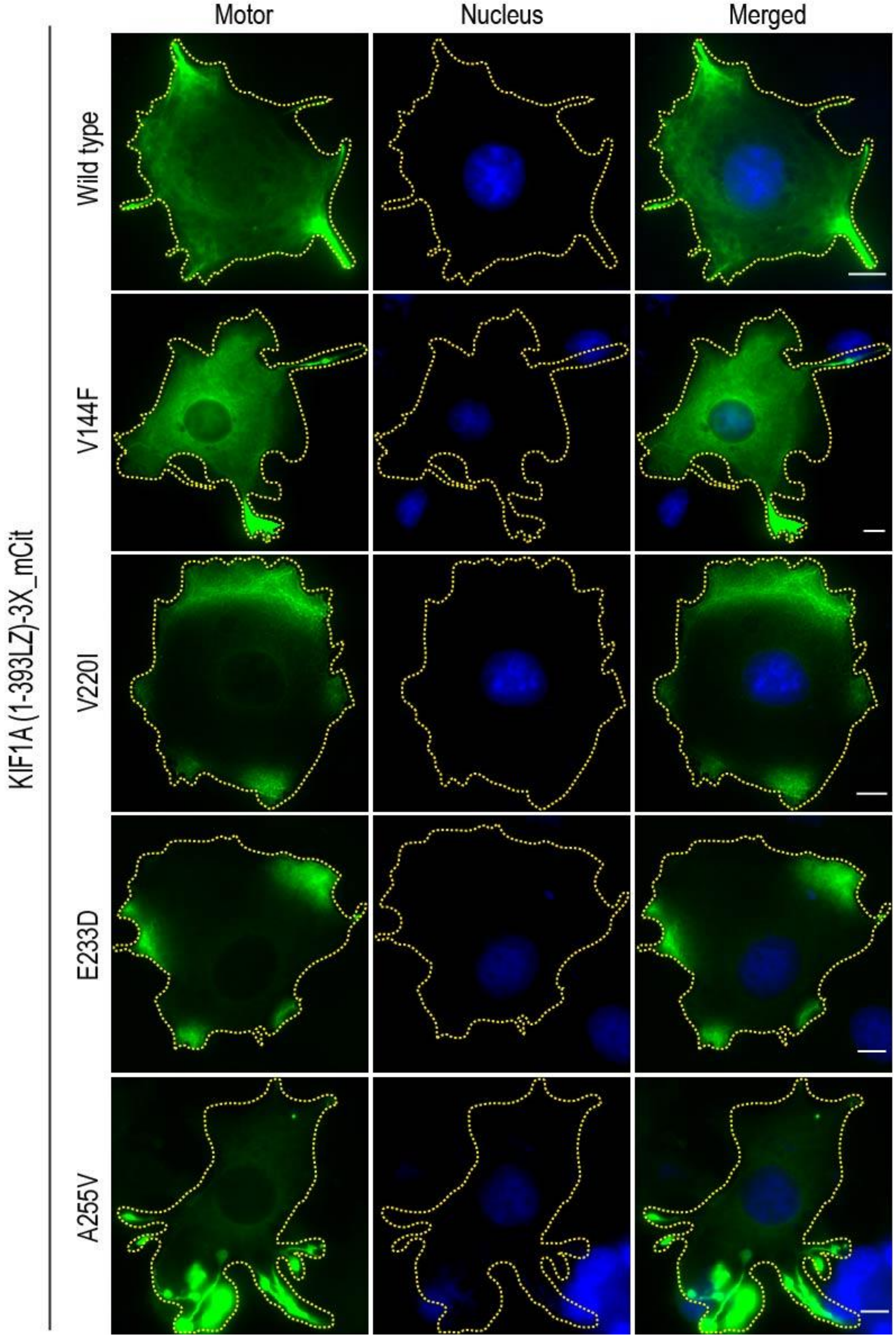


**Figure S1:** Expression of WT and V144F, V220I, E233D, and A255V mutants showing peripheral accumulation comparable to WT motor. The green channel represents the motor, and the blue channel shows DAPI staining. Scale Bar: 10µm.

Figure S2

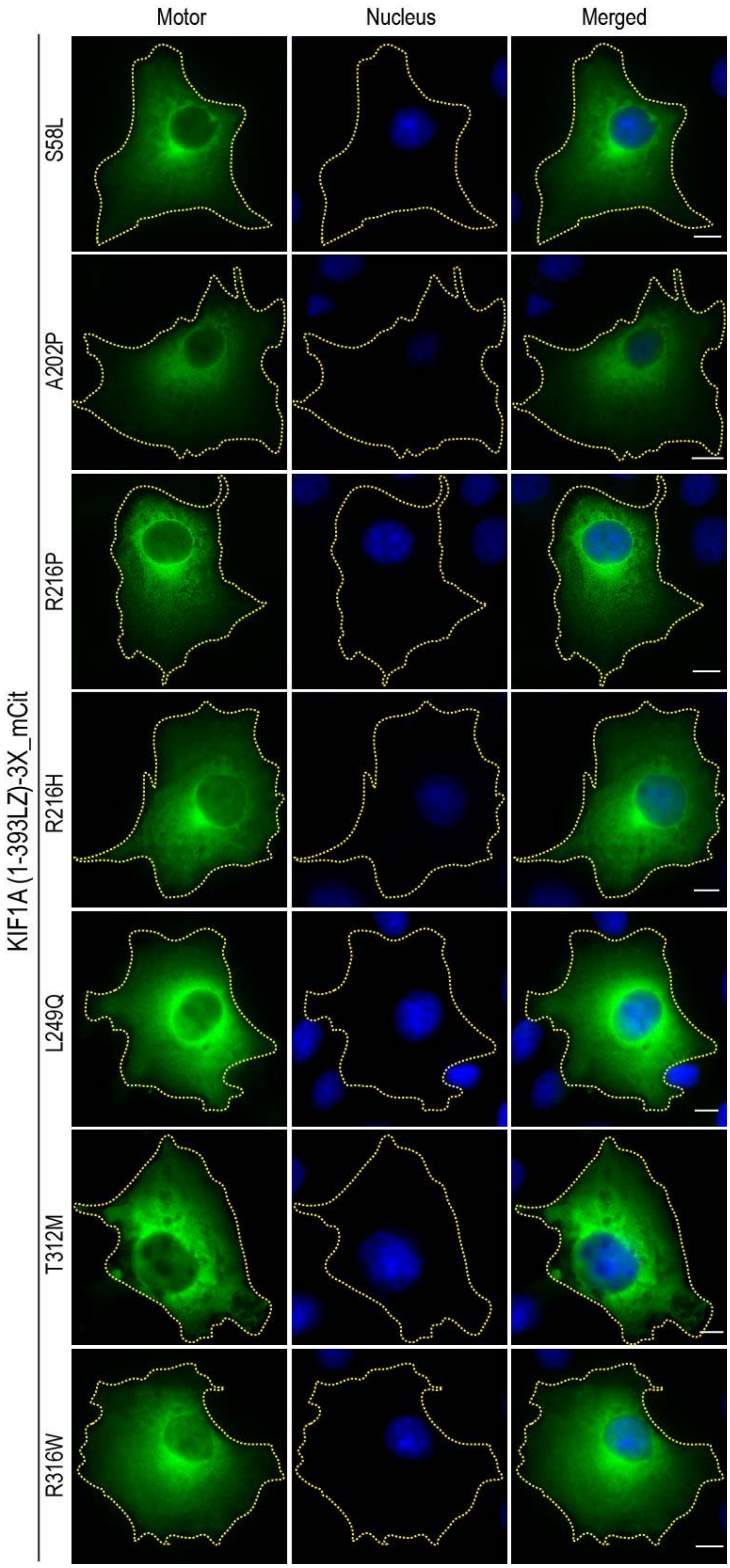
Motor
Nucleus
Merged
KIF1A (1-393LZ)-3X_mCit
S58L
A202P
R216P
R216H
L249Q
T312M
R316W

Figure S2: TIRF images of S58L, A202P, R216P, R216H, L249Q, R316W, and R350G mutants showing diffused cytoplasmic phenotype in COS-7 assay. Motors are seen in the green channel, and the blue channel is nuclear staining by DAPI. Scale Bar: 10µm.

Figure S3

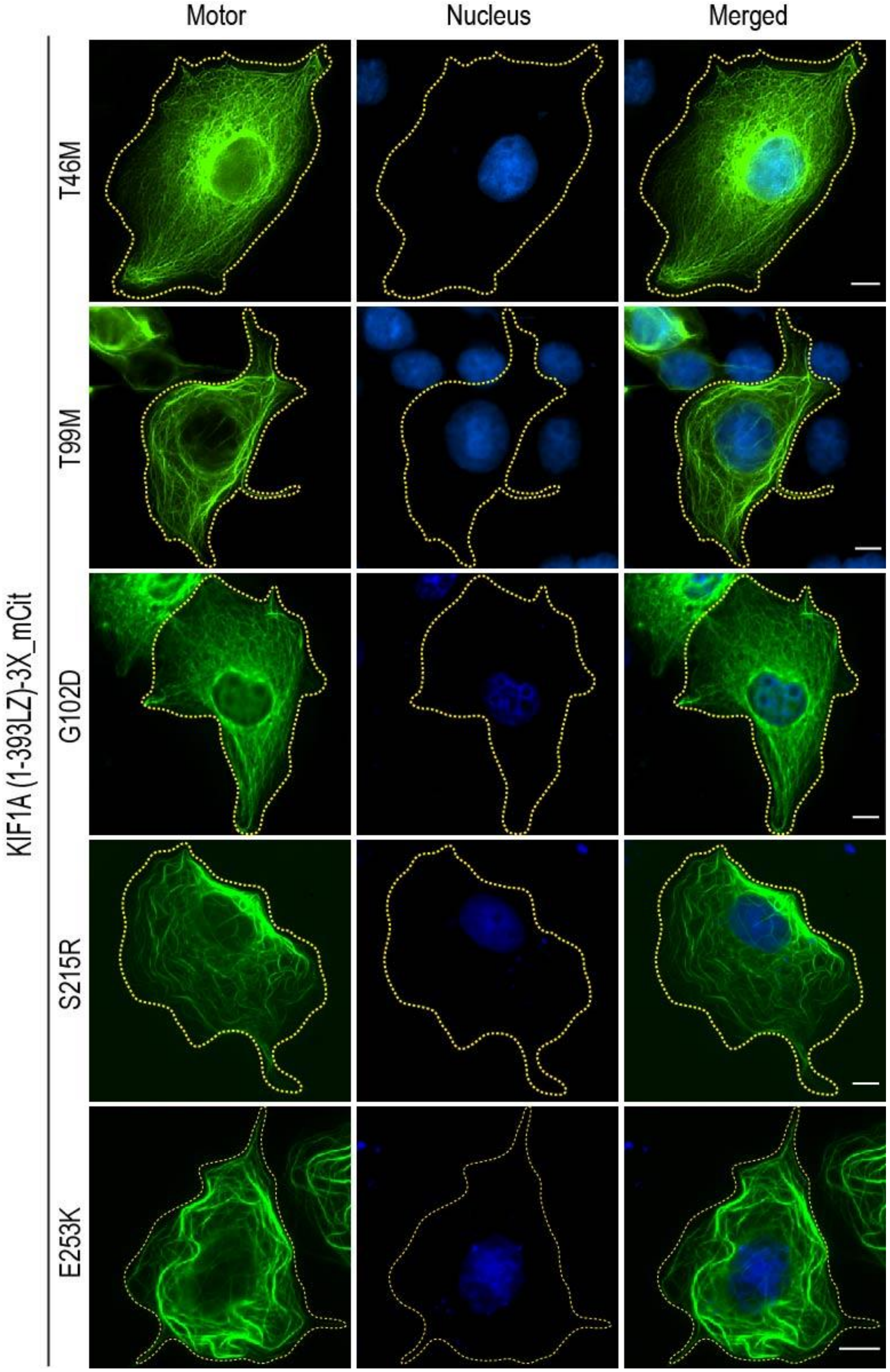


Figure S3: Expression of T46M, T99M, G102D, S215R, and E253K mutants in COS-7 cells showing decoration of microtubules. The green channel represents the motor, and blue is DAPI staining for the nucleus. Scale Bar: 10µm.

Figure S4

A

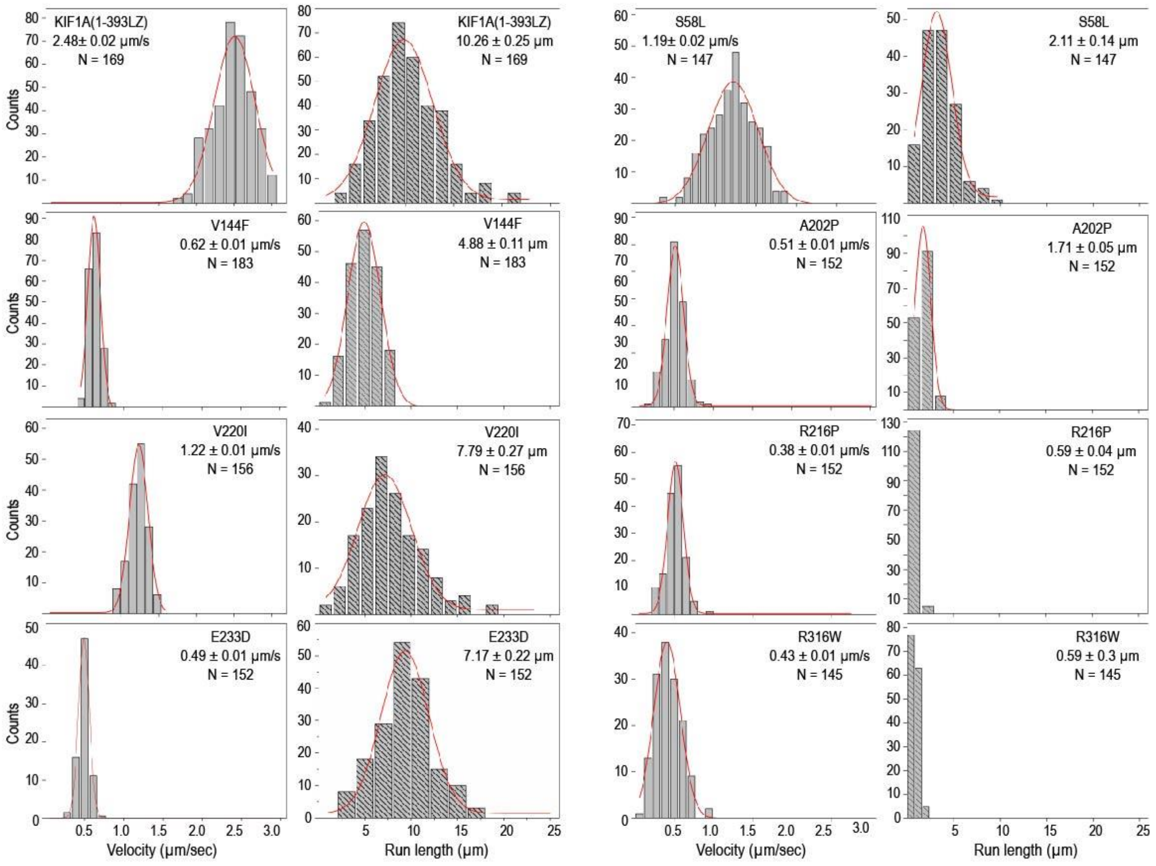


B

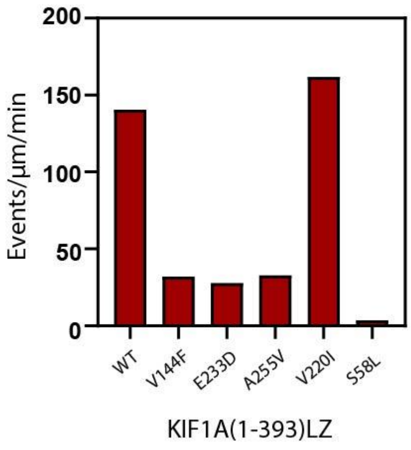


Figure S4: A. Velocities from motility assay of WT and motile mutants were plotted in the form of histograms and fitted to a Gaussian distribution, mutant name, velocity, or run length along with tracked population of motors (N) is indicated at the top corner of the histograms as mean±SEM. Data is presented from three independent experiments. B. Landing rate for WT and mutant motors calculated from motility events observed on ~150 μm microtubule length for each mutant.

Figure S5

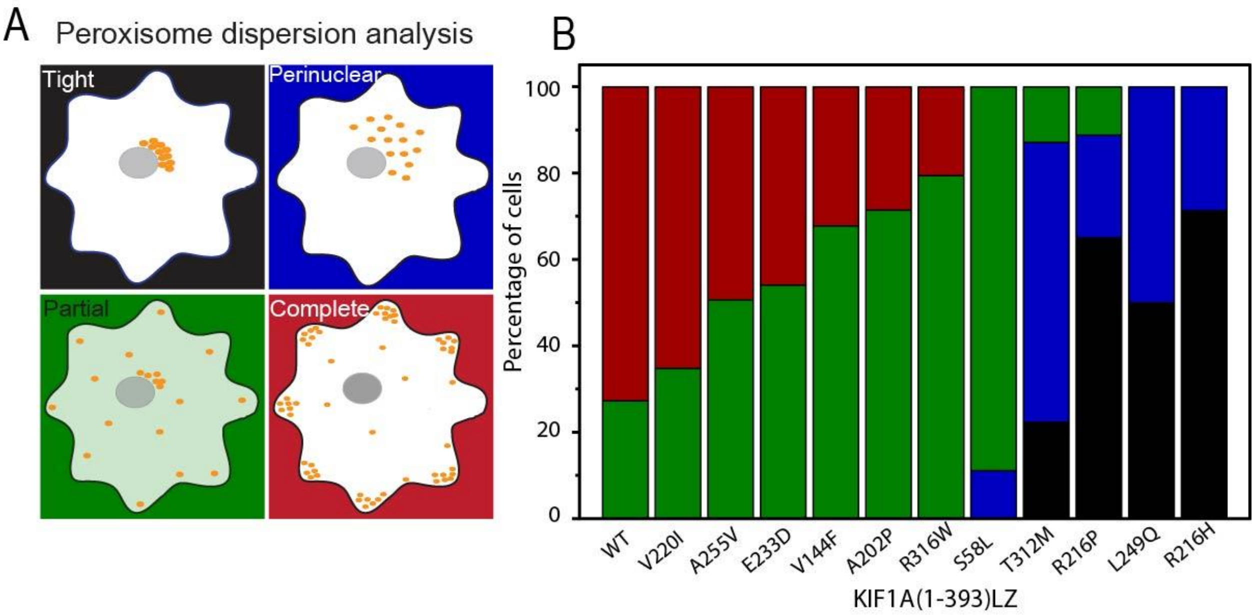


Figure S5: **A.** Graphical representation of four cellular phenotypes of peroxisome dispersion, peripheral (red), partial(green), perinuclear (blue), and tight(black), **B.** Bar graph representing percentage cells dispersing the peroxisome to particular phenotype mentioned in (A) for WT and mutant motors.

Figure S6

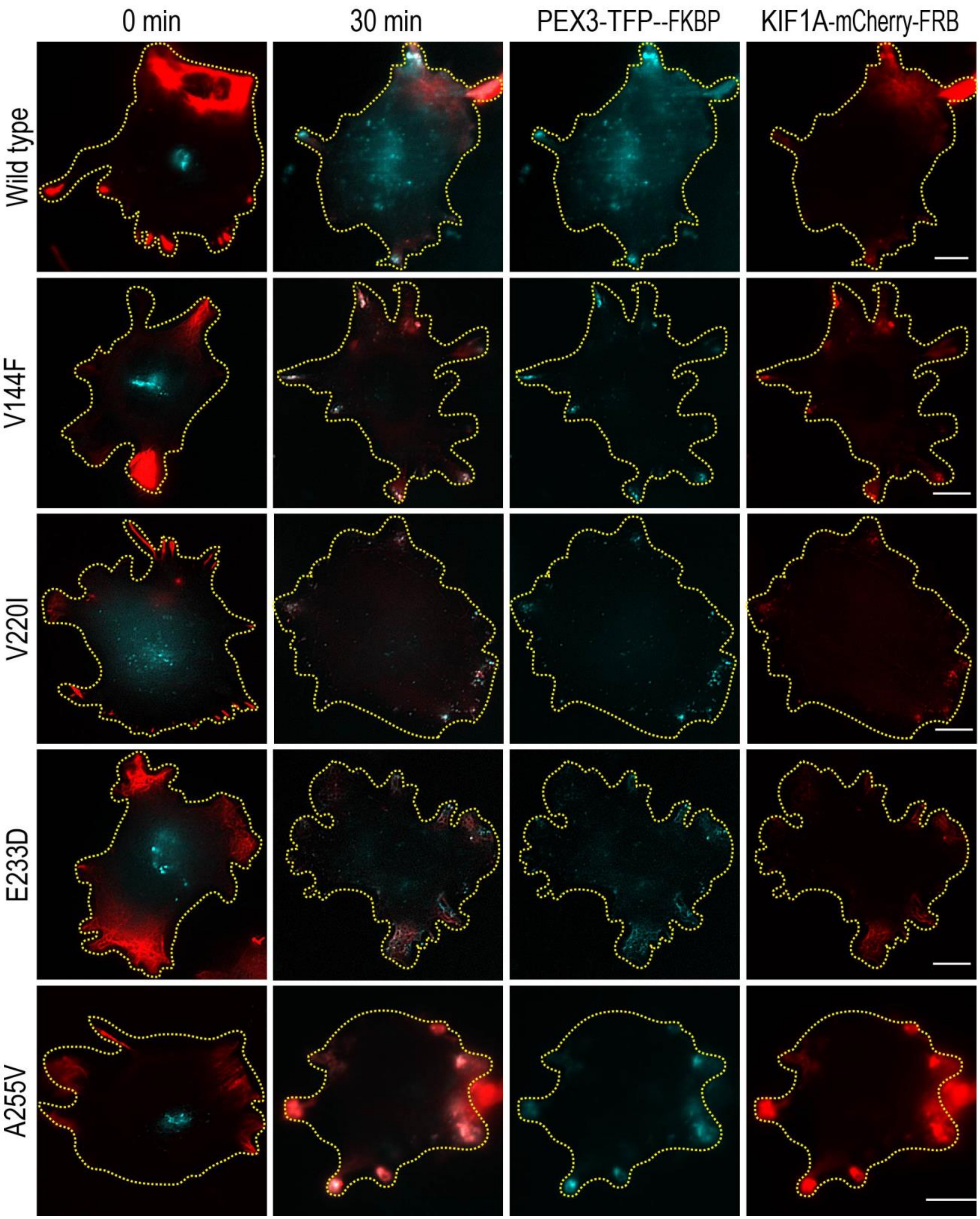

0 min
30 min
PEX3-TFP--FKBP
KIF1A-mCherry-FRB
Wild type
V144F
V220I
E233D
A255V

Figure S6: TIRF images of COS-7 cells expressing WT and V144F, V220I, E233D, and A255V mutants tagged with mCherry and peroxisome marked with a teal fluorescent protein; the first panel shows merged images of unloaded motor (red) with peroxisome (cyan) in the perinuclear region before rapamycin treatment. The second panel shows the motor loaded on peroxisomes and transported to the periphery. The last two panels are individual channels displaying PEX3-TFP-FKBP (in cyan) and Motor-mCherry-FRB (in red), respectively. Scale bar: 10μm.

Figure S7

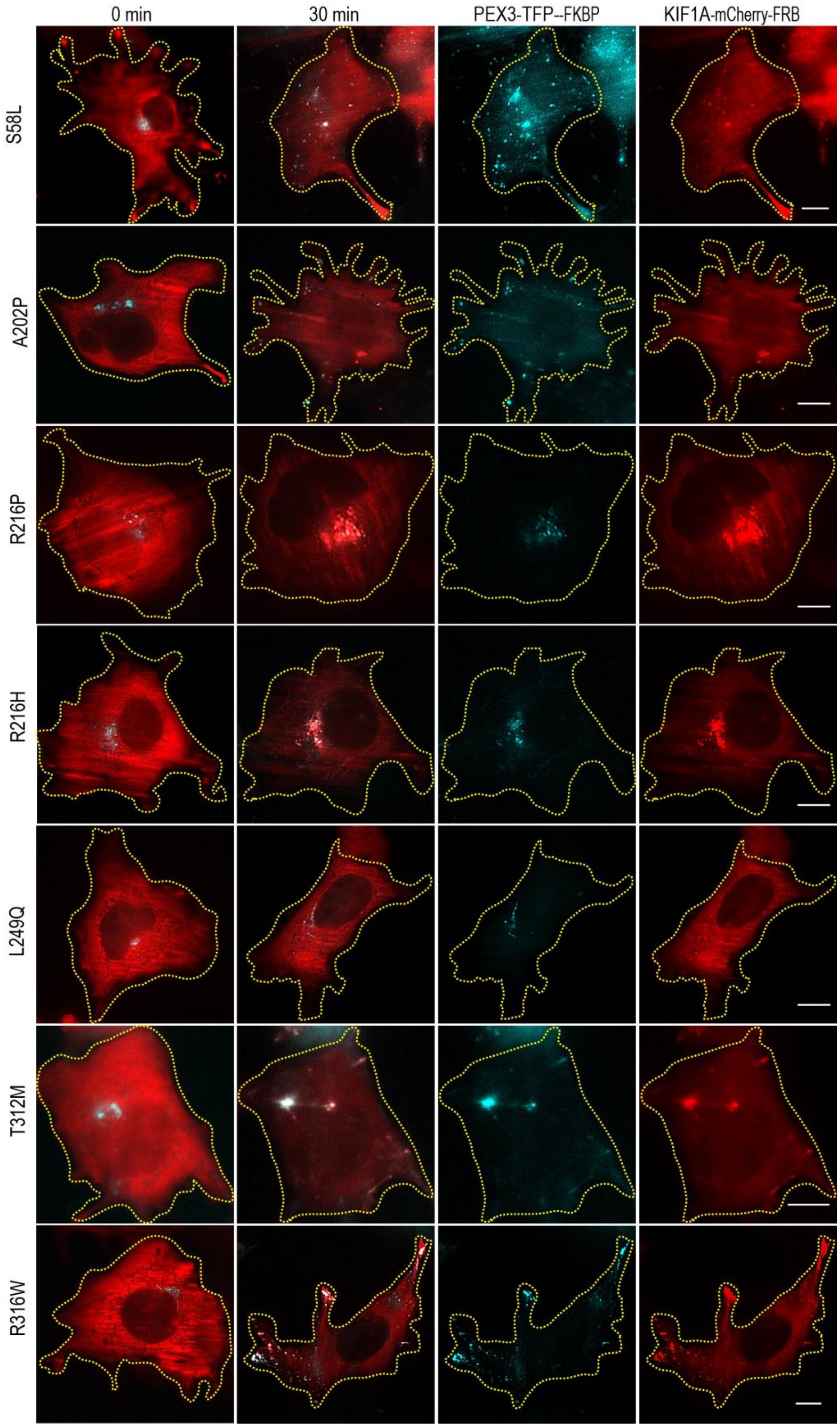

0 min
30 min
PEX3-TFP--FKBP
KIF1A-mCherry-FRB
S58L
A202P
R216P
R216H
L249Q
T312M
R316W

Figure S7: TIRF images showing unloaded diffused motor (red) and tight peroxisome phenotype (cyan) in the first panel after 0 min of rapamycin addition. The second panel shows a merged image of the motor loaded on the peroxisome after 30 min of rapamycin treatment showing dispersion by S58L, A202P, R216P, R216H, L249Q, T312M, and R316W mutants. The third and fourth panels are separate red and cyan channels for images in the second panel. Scale bar: 10µm.

Figure S8

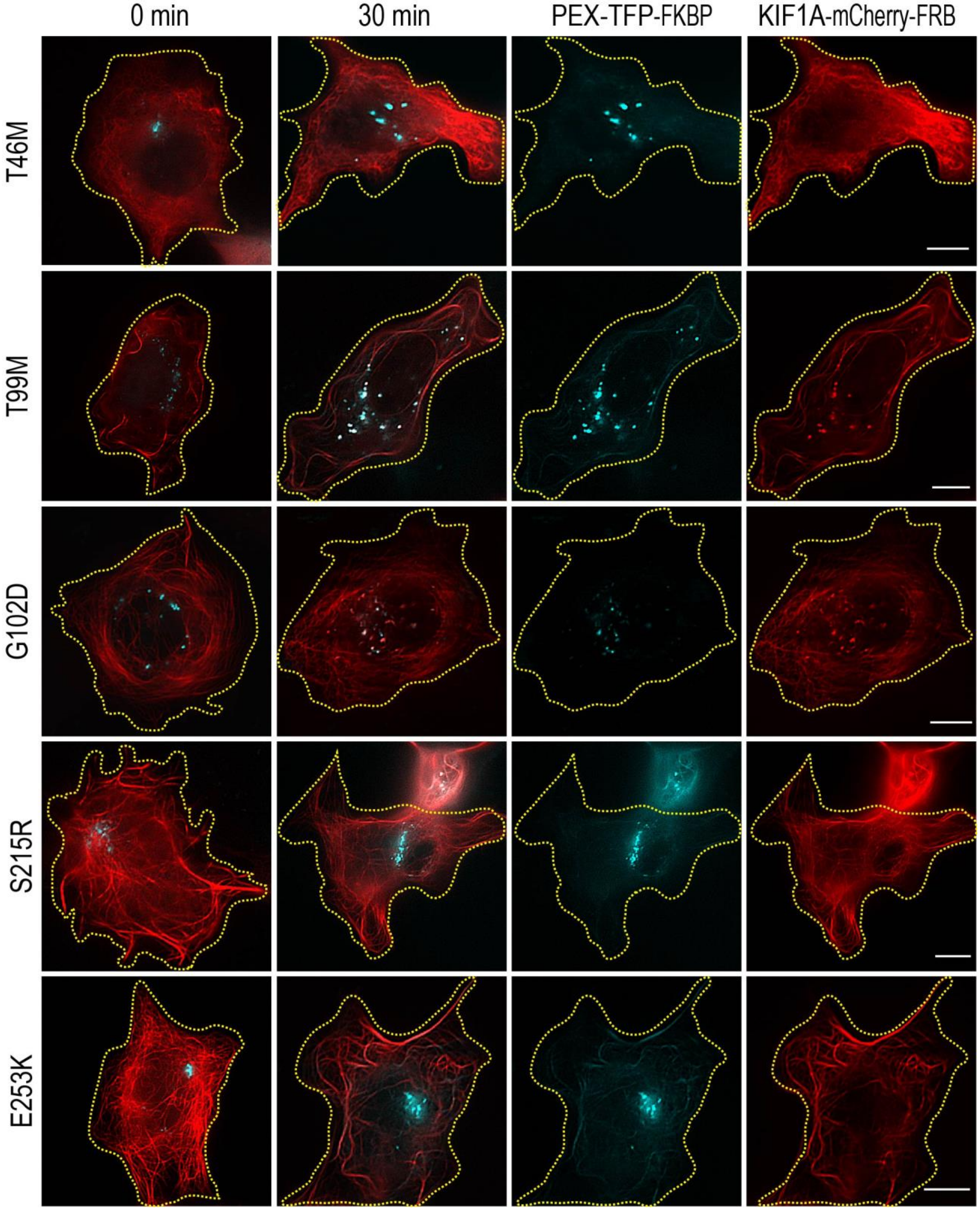

0 min
30 min
PEX-TFP-FKBP
KIF1A-mCherry-FRB
T46M
T99M
G102D
S215R
E253K

Figure S8: The first panel shows peroxisome (cyan) localization at 0 min of rapamycin treatment when MT-bound mutant motor (red) is not loaded on peroxisomes. The second panel displays a merged image of mutant motor and peroxisome dispersion after 30 min of rapamycin treatment. The next two panels show separate channels (red and cyan) for the merged image in the second panel. Scale bar: 10µm.

Figure S9

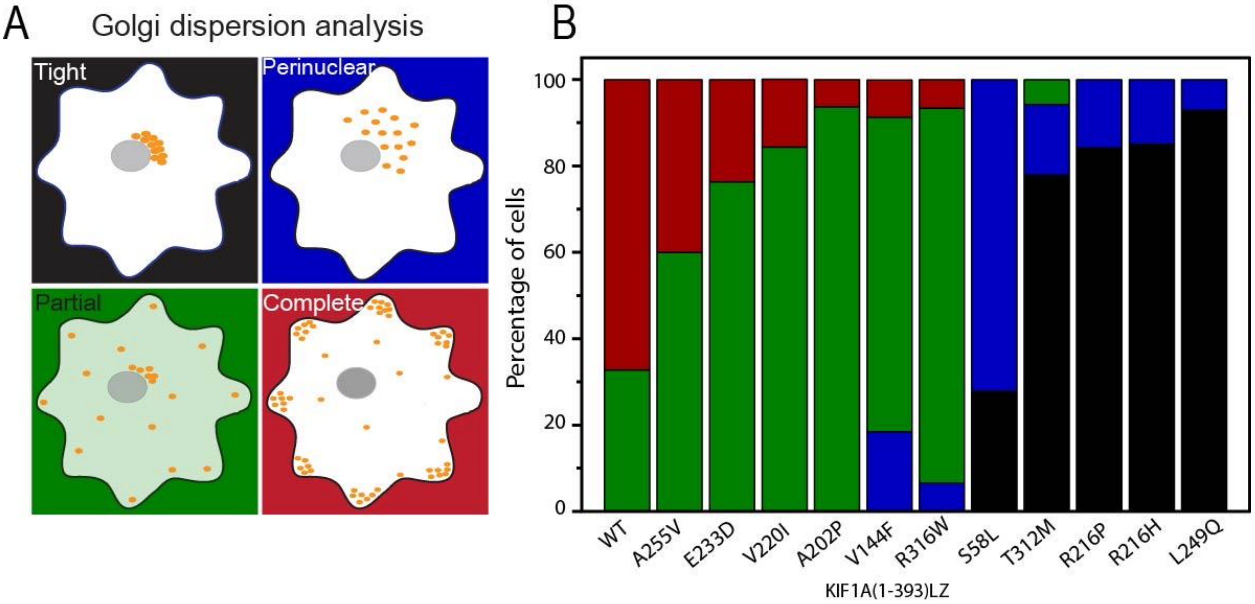


Figure S9: **A.** Cartoon representing four types of Golgi distribution by KIF1A mutant motors after rapamycin treatment, tight, perinuclear, partial, and peripheral. **B.** Percentage cells displaying particular phenotype of Golgi dispersion by WT and KIF1A mutants.

Figure S10

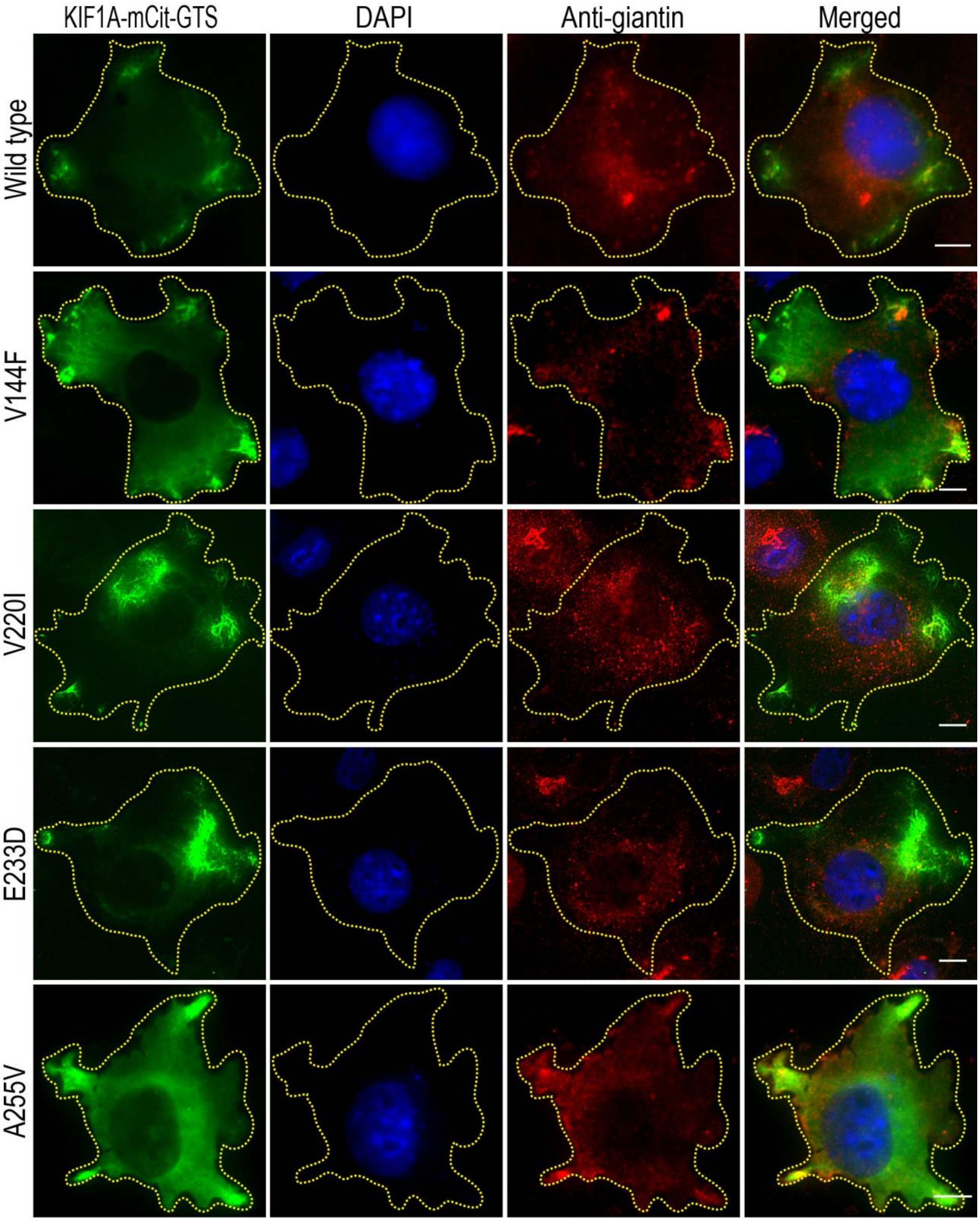

KIF1A-mCit-GTS
DAPI
Anti-giantin
Merged
Wild type
V144F
V220I
E233D
A255V

Figure S10: Green channel shows the TIRF images of the KIF1A mutant motor tagged with GTS sequence expressed in COS-7 cells. The blue channel is nuclear staining with DAPI dye, and the red channel displays Golgi stained with an anti-giantin antibody. The fourth panel shows a merged image for WT and each mutant (V144F, V220I, E233D, and A255V).

Figure S11

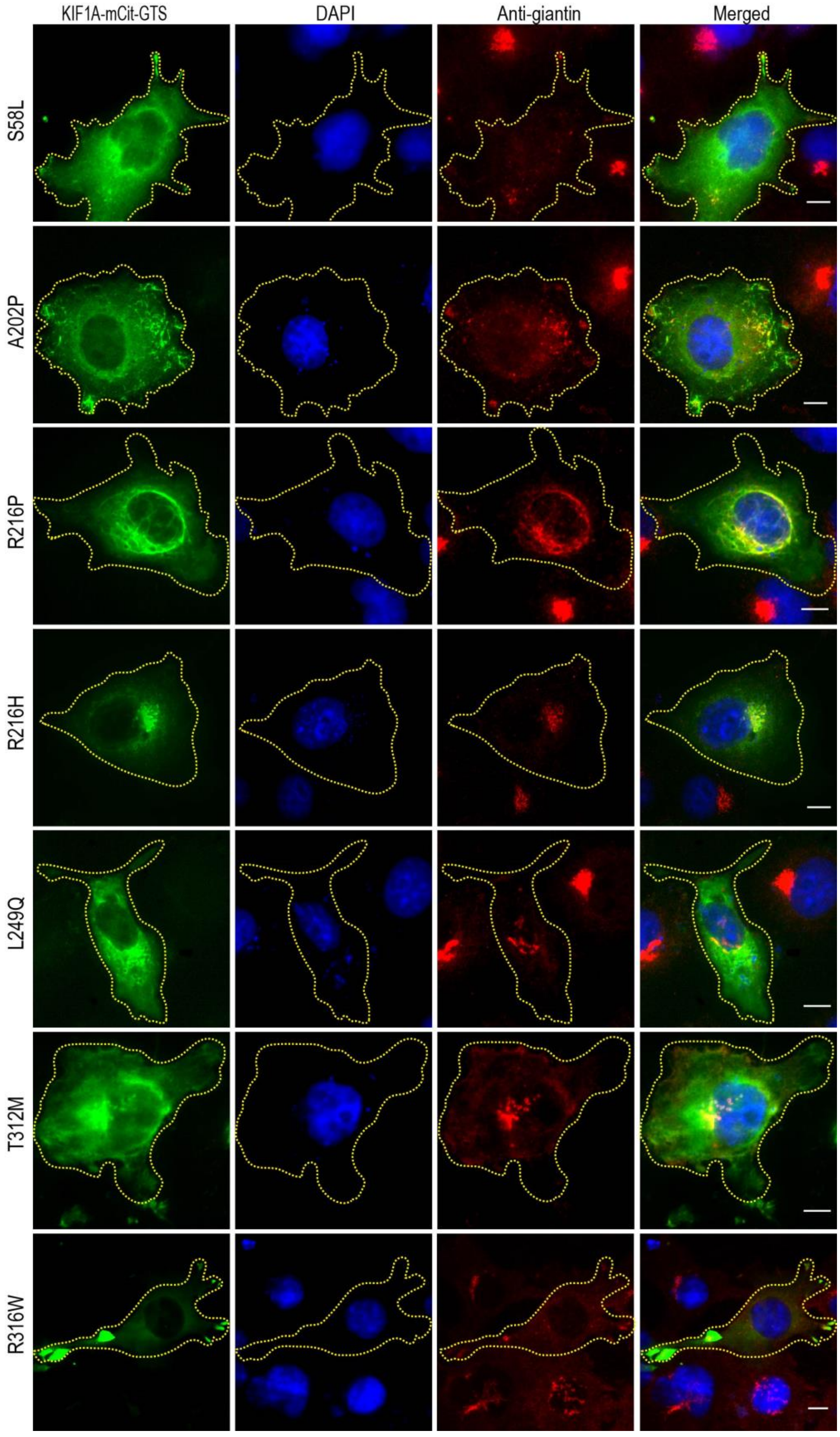
KIF1A-mCit-GTS
DAPI
Anti-giantin
Merged
S58L
A202P
R216P
R216H
L249Q
T312M
R316W

Figure S11: The green channel shows a cytoplasmic KIF1A mutant motor with a GTS sequence in COS-7 cells. The blue channel is DAPI staining for the nucleus, and the red staining with anti-giantin antibody marks Golgi. Fourth panel shows merged images for S58L, A202P, R216P, R216H, L249Q, T312M and R316W mutants.

Figure S12

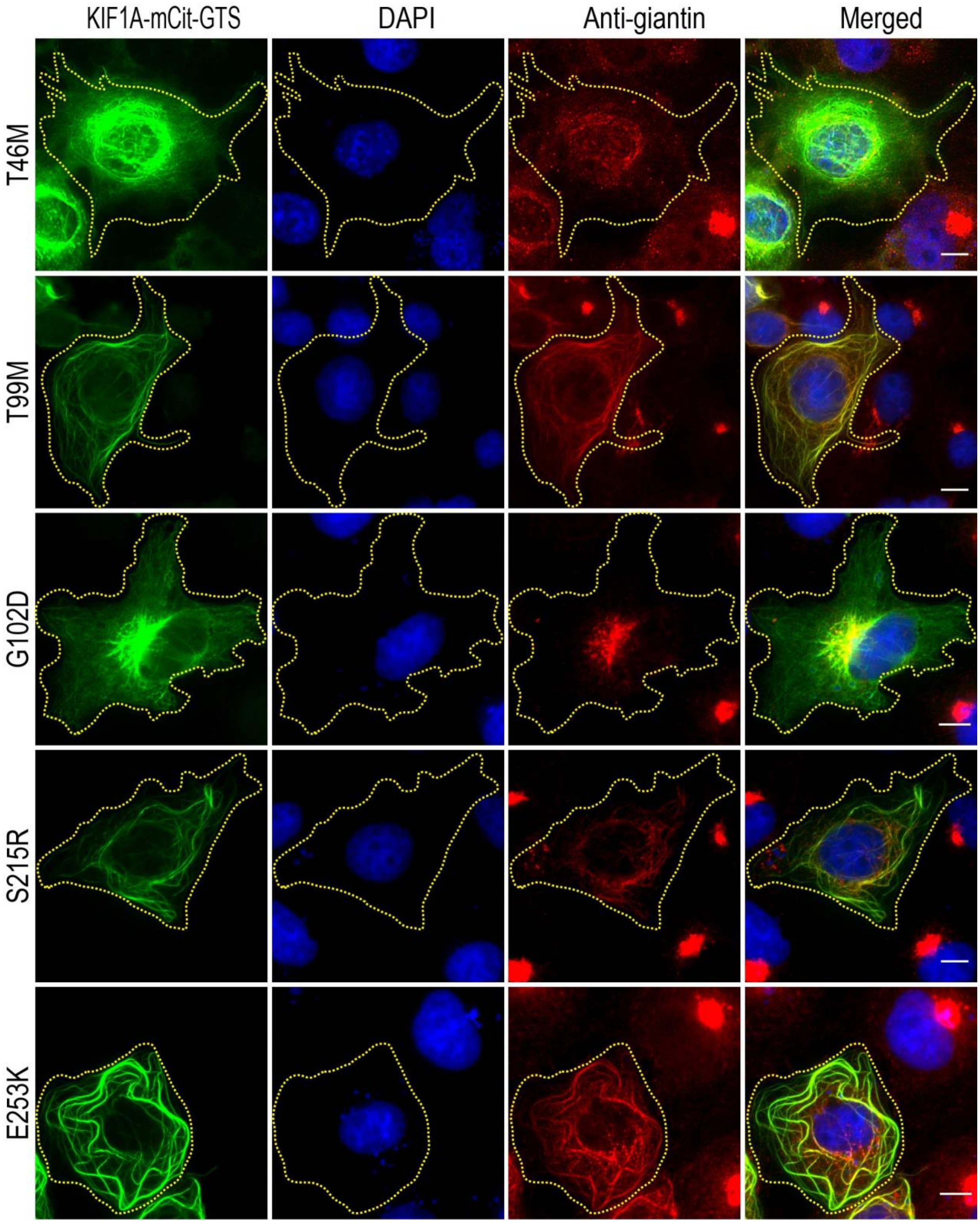
KIF1A-mCit-GTS
DAPI
Anti-giantin
Merged
T46M
T99M
G102D
S215R
E253K

Figure S12: The first panel shows MT-bound KIF1A mutant motor (green) tagged with GTS sequence in COS-7 cells. DAPI staining was used for visualizing the nucleus (blue), and anti-giantin antibody marks Golgi (red). The fourth panel displays merged images for all three channels showing Golgi dispersion by T46M, T99M, G102D, S215R, and E253K mutants

### Movie legends

Movie 1: Microtubule gliding by constitutively active KIF1A WT motor coated on GFP-nanobody.

Movie 2: Microtubule gliding by constitutively active KIF1A motor with V144F mutation.

Movie 3: Microtubule gliding by constitutively active KIF1A motor with E233D mutation.

Movie 4: Microtubule gliding assay with constitutively active KIF1A motor containing V220I mutation.

Movie 5: Microtubule gliding by constitutively active KIF1A motor with A255V mutation. Arrowheads denote the spiralling of microtubules while gliding on the mutant motors.

Movie 6: Movie demonstrating intracellular vesicle transport by full-length KIF1AWT motor in COS-7 cells.

Movie 7: Intracellular vesicle transport by full-length KIF1A(V144F) mutant motor in COS-7 cells.
